## Supplementary Material for "Reconstructing Voice Identity from Noninvasive Auditory Cortex Recordings"

### Supplementary Information

- Figure 1-S1. Projections of the DNN-derived Voice Latent Space (VLS).
- Figure 2-S1. Brain activity in response to voice measured by fMRI.
- Figure 2-S2. Denoising of the fMRI BOLD responses.
- Figure 2-S3. Extended predicted brain encoding results with state-of-the-art models.
- Supplementary Table 1. Architecture of the VAE network.
- Supplementary Table 2. Assessing significance of brain encoding performance with LIN features.
- Supplementary Table 3. Assessing significance of brain encoding performance with VLS features.
- Supplementary Table 4. Comparing the performance of brain encoding models.
- Supplementary Table 5. Comparing the performance of brain encoding ROIs.
- Supplementary Table 6. Assessing significance of the RSA brain-model correlation.
- Supplementary Table 7. Comparing the performance of the RSA models.
- Supplementary Table 8. Assessing the significance of speaker gender decoding performance using VLS and LIN models based on voxel activity.
- Supplementary Table 9. Assessing the significance of speaker age decoding performance using VLS and LIN models based on voxel activity.
- Supplementary Table 10. Assessing the significance of speaker identity decoding performance using VLS and LIN models based on voxel activity.
- Supplementary Table 11. Comparing the performance of the models decoding speaker identity-related information.
- Supplementary Table 12. Comparing the performance of the models decoding speaker identity-related information by ROI.

- Supplementary Table 13. Assessing the significance of the speaker gender discrimination task.
- Supplementary Table 14. Assessing the significance of the speaker age discrimination task.
- Supplementary Table 15. Assessing the significance of the speaker discrimination task.
- Supplementary Table 16. Comparing human listeners' performance in discriminating speaker identity-related information decoded with VLS versus LIN.
- Supplementary Table 17. Comparing the performance of the human listeners at discriminating speaker identity-related information by ROI.
- Supplementary Audio 1. Voice latent space interpolation.
- Supplementary Audio 2. Brain- and spectrogram -based voice reconstructions.

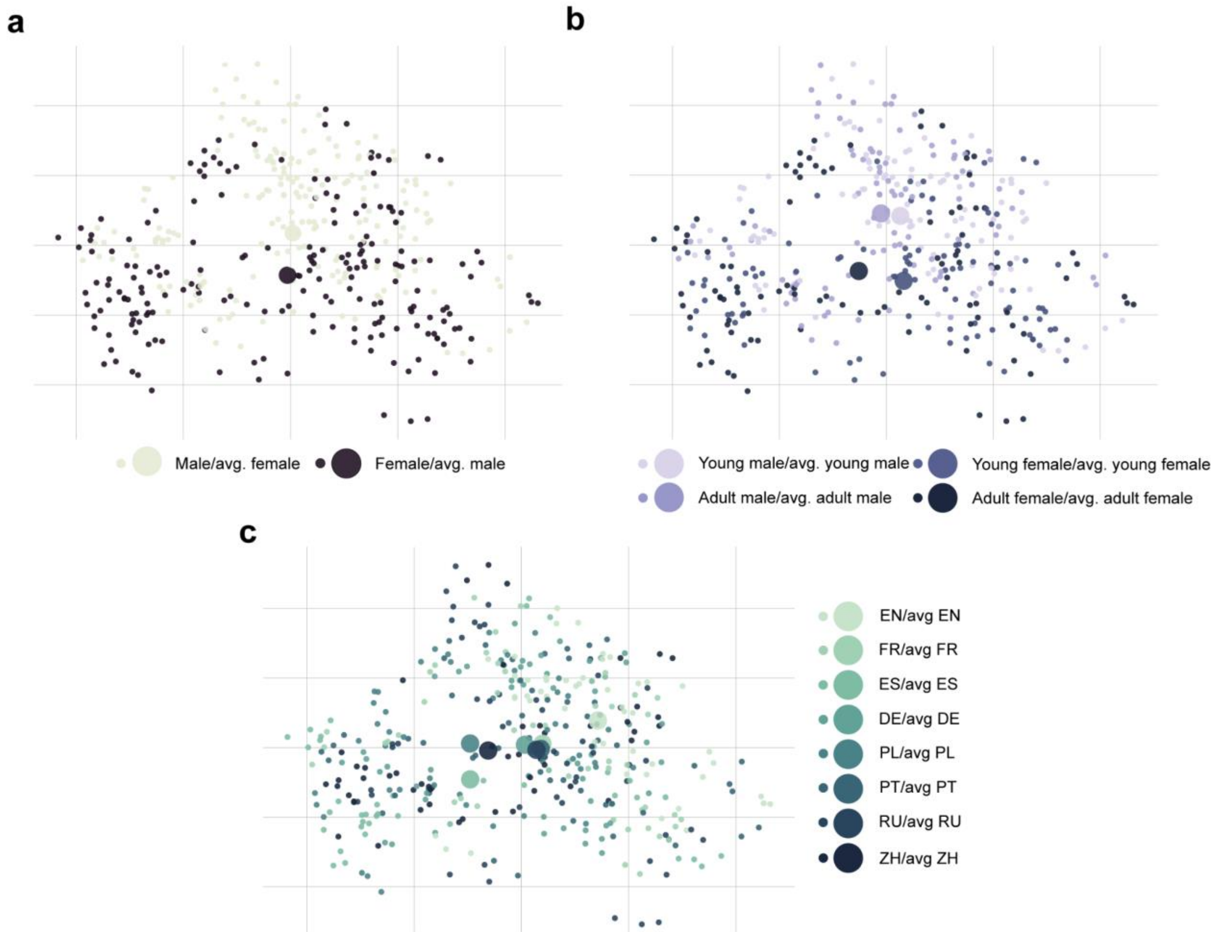


**Figure 1-S1. Projections of the DNN-derived Voice Latent Space (VLS).** Distribution of the 405 speaker identities along the first 2 principal components of the VLS coordinates from all sounds, averaged by speaker identity. Each disk represents a speaker's identity colored by either gender (as in Fig. 1b), age, or language. **a,** Large disks represent the average of all male (black) or female (gray) speaker coordinates. ANOVAs on the first two components: PC1: $F(1, 405)=0.10$, $p=.74$; PC2: $F(1, 405)=11.00$, $p<.001$. **b,** Same for speaker age. ANOVAs on first two components: PC1: $F(1, 405)=4.12$, $p<.01$; PC2: $F(1, 405)=3.99$, $p<.01$. **c,** Same for speaker language. ANOVAs on the first two components: PC1: $F(1, 405)=8.46$, $p<.0001$; PC2: $F(1, 405)=6.09$, $p<.0001$.

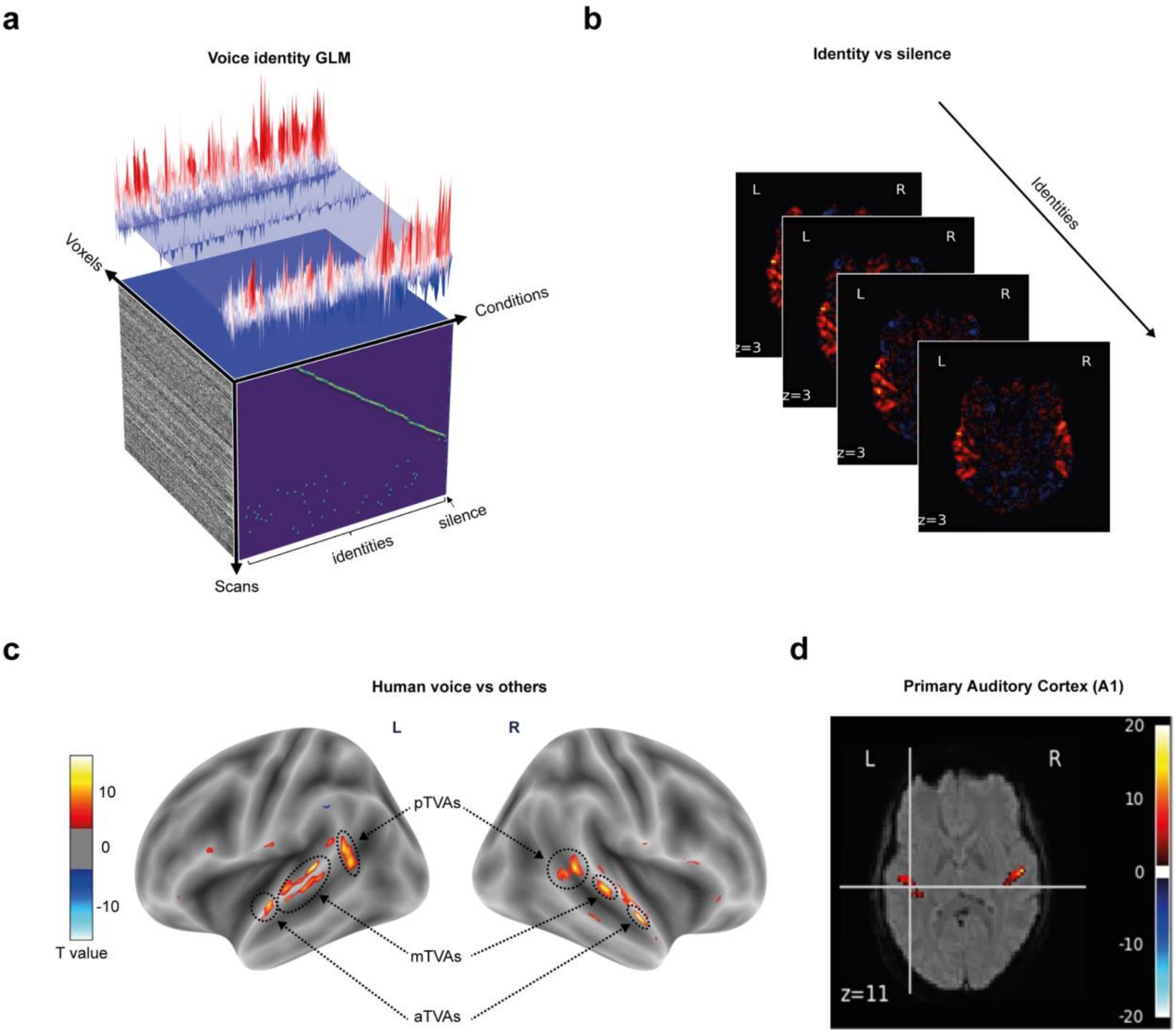


**Figure 2-S1. Brain activity in response to voice measured by fMRI. a,** A GLM is used to model fMRI activity in response to each speaker's identity. **b,** The fMRI activity in response to each speaker's identity is mapped into dedicated voxel maps by contrasting the speaker's identity with the silence, resulting in ~135 voxel maps. **c,** The voice-sensitive ROIs used for subsequent analyses, identified in each participant via an independent Voice Localizer: the anterior, middle, and posterior Temporal Voice Areas (TVAs). **d,** The Primary Auditory Cortex (A1) is defined as the intersection between a probabilistic map of Heschl's gyri and the sound vs silence contrast map.

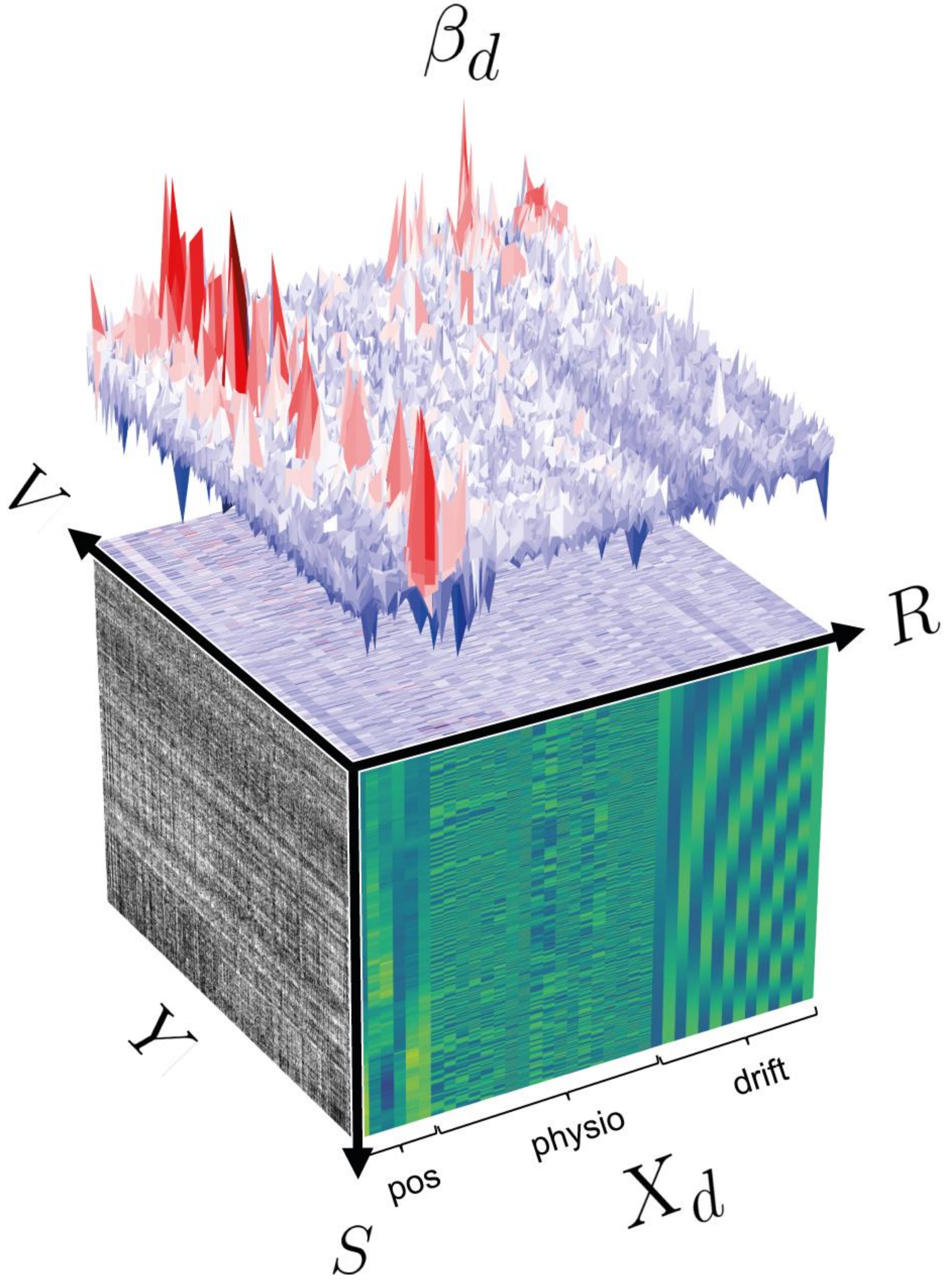


**Figure 2-S2. Denoising of the fMRI BOLD responses.** A general linear model (GLM) was fit to regress out the noise by predicting $Y$ from a "denoising" design matrix $X_d$ , composed of $R = 38$ regressors of nuisance 6 head motion parameters (3 variable for the translations, 3 variables for the rotations); 18 'RETROICOR' regressors (Glover et al., 2000) using the *TAPAS PhysIO* package (Kasper et al., 2017) with the hyperparameters set as specified in (Snoek et al., 2021); 13 regressors modeling slow artifactual trends (sines and cosines, cutoff frequency of the high-pass filter = 0.01 Hz); an intercept. The design matrix was convolved with an hemodynamic response function (HRF) with a peak at 6s sec and an undershoot at 16s sec (Glover et al., 1999), we note the convolved design matrix as $X_d \in R^{S \times R}$ where $S$ = number of scans. The “denoise” GLM’s parameters $\beta_d \in R^{R \times V}$ were optimized to minimize the amplitude of the residual $\beta_d = argmin_{\beta \in R^{R \times V}} \ || \ Y \ - X_d \ \beta \ ||^2$, where $V$ = number of voxels. The *denoised* BOLD signal $Y_d$ was then obtained from the original one according to $Y_d = Y - (X_d \ \beta_d)$.

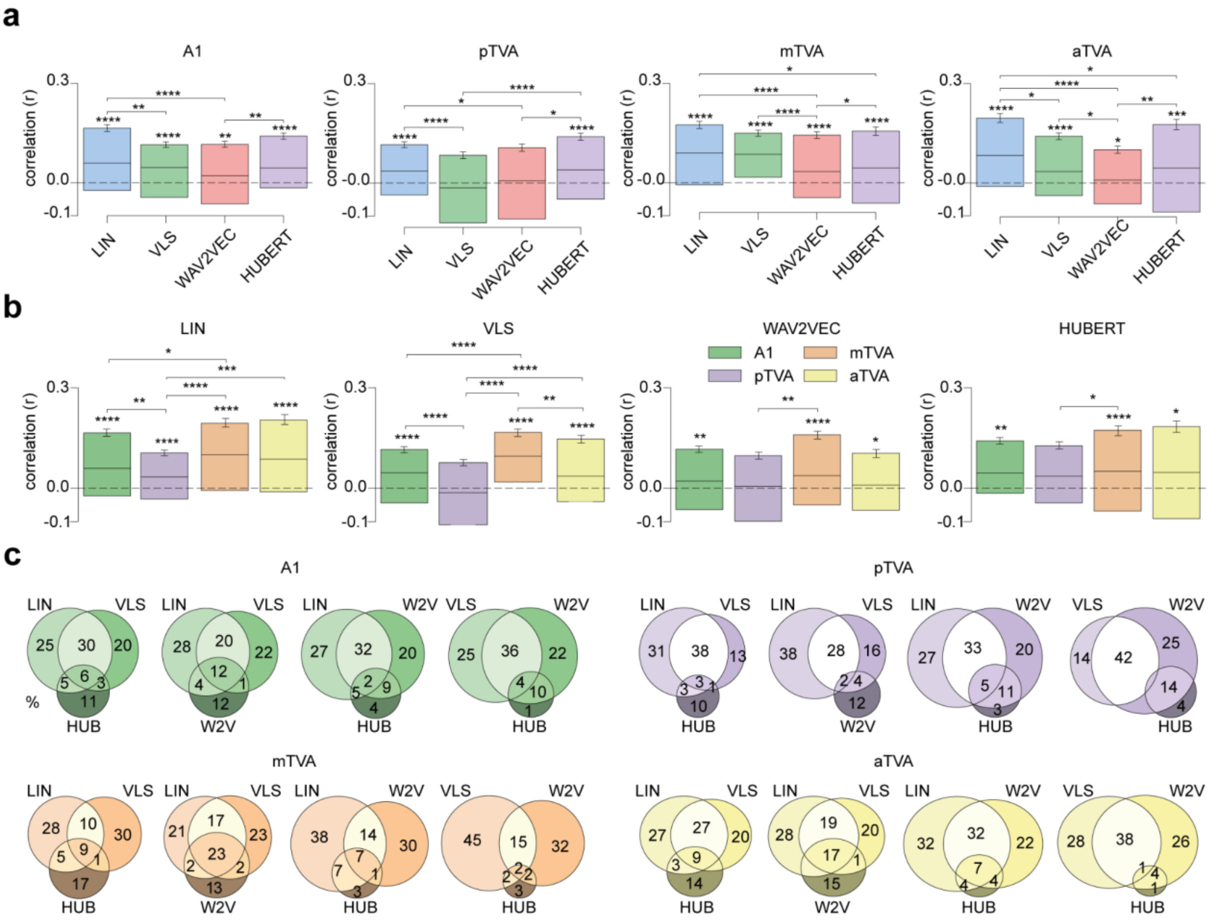


**Figure 2-S3. Extended predicted brain encoding results with state-of-the-art models. a,** Encoding results with Wav2Vec and HuBERT. For each region of interest, the model's performance was assessed using the Pearson correlation score between the true and the predicted responses of each voxel on the held-out speaker identities. Pearson’s correlation coefficients were computed for each voxel on the speakers’ axis and then averaged across hemispheres and participants.. Error bars indicate the standard error of the mean (s.e.m) across voxels. *p < 0.05; **p < 0.01; **p < 0.001; ****p < 0.0001. **b,** Same results grouped by model. **c,** Venn diagrams of the number of voxels for each ROI and triplet of models. For each ROI and each voxel, we checked whether the test correlation was higher than the median of all participant correlations (intersection circle), and if not, which model yielded the highest correlation (left or right circles).

| Name | Layer | #Filters | Filter size | Stride | Activation |
|---|---|---|---|---|---|
| Encoder | Conv2D + BN2D | 64 | 6x3 | 2x2 | ReLU |
| | Conv2D + BN2D | 128 | 6x2 | 2x2 | ReLU |
| | Conv2D + BN2D | 256 | 6x2 | 2x1 | ReLU |
| | Conv2D + BN2D | 512 | 6x2 | 2x1 | ReLU |
| | Conv2D | 7 | 6x2 | 1x1 | - |
| Bottleneck | FC | 256 | - | - | - |
| Decoder | ConvTrans2D + BN2D | 512 | 27x3 | 1x1 | ReLU |
| | ConvTrans2D + BN2D | 256 | 4x2 | 2x1 | ReLU |
| | ConvTrans2D + BN2D | 128 | 4x2 | 2x1 | ReLU |
| | ConvTrans2D + BN2D | 64 | 4x2 | 2x2 | ReLU |
| | ConvTrans2D | 1 | 4x2 | 2x2 | - |
| Batch size | 64 | | | | |
| Loss function | MSE + KL divergence | | | | |
| Optimizer | Adam, learning rate = 0.00005 | | | | |
| | betas = (0.5, 0.999) | | | | |

**Supplementary Table 1. Architecture of the VAE network.** The architecture of the VAE consists of 15 layers with an intermediate hidden representation of 128 neurons that will stand for the VLS. The Encoder network ($Enc$; 7 layers) learns to map an input, $s$ (a spectrogram of a sound), onto the (128-dimensional) VLS, while the Decoder ($Dec$; 7 layers) aims at reconstructing the spectrogram $s$ from $z$. The learning objective of the full model is to make the output spectrogram $Dec(Enc(s))$ as close as possible to the original one $s$. The model was trained until convergence (approximately 1000 epochs). Hyperparameter search was conducted to determine the suitable learning rate. BN: batch normalization; FC: fully connected; ReLU: Rectified Linear Unit.

| Subject | ROI | Correlation | s.e.m. | T | dof | p-val | unc. sig. | CI95% | cohen-d | BF10 | power |
|---|---|---|---|---|---|---|---|---|---|---|---|
| s1 | LA1 | 0.13 ± 0.15 | 0.03 | 4.78E+00 | 32 | 1.91E-05 | **** | [0.08, inf] | 8.30E-01 | 1.22E+03 | 1.00 |
| | RA1 | 0.21 ± 0.14 | 0.03 | 8.08E+00 | 32 | 1.57E-09 | **** | [0.16, inf] | 1.41E+00 | 7.74E+06 | 1.00 |
| | LmTVA | 0.32 ± 0.13 | 0.02 | 1.34E+01 | 32 | 5.25E-15 | **** | [0.28, inf] | 2.34E+00 | 1.27E+12 | 1.00 |
| | RmTVA | 0.16 ± 0.07 | 0.01 | 1.11E+01 | 26 | 1.21E-11 | **** | [0.13, inf] | 2.13E+00 | 7.53E+08 | 1.00 |
| | LpTVA | 0.07 ± 0.13 | 0.02 | 3.15E+00 | 32 | 1.76E-03 | ** | [0.03, inf] | 5.50E-01 | 2.14E+01 | 0.92 |
| | RpTVA | 0.04 ± 0.08 | 0.02 | 2.56E+00 | 31 | 7.82E-03 | ** | [0.01, inf] | 4.50E-01 | 6.05E+00 | 0.80 |
| | LaTVA | 0.27 ± 0.15 | 0.03 | 1.00E+01 | 30 | 2.30E-11 | **** | [0.23, inf] | 1.80E+00 | 4.20E+08 | 1.00 |
| | RaTVA | 0.11 ± 0.10 | 0.02 | 5.26E+00 | 25 | 9.42E-06 | **** | [0.07, inf] | 1.03E+00 | 2.42E+03 | 1.00 |
| | A1 | 0.17 ± 0.15 | 0.02 | 8.80E+00 | 65 | 5.58E-13 | **** | [0.14, inf] | 1.08E+00 | 1.48E+10 | 1.00 |
| | mTVA | 0.25 ± 0.14 | 0.02 | 1.38E+01 | 59 | 1.71E-20 | **** | [0.22, inf] | 1.79E+00 | 2.85E+17 | 1.00 |
| | pTVA | 0.06 ± 0.11 | 0.01 | 4.02E+00 | 64 | 7.84E-05 | **** | [0.03, inf] | 5.00E-01 | 2.81E+02 | 0.99 |
| | aTVA | 0.20 ± 0.15 | 0.02 | 9.63E+00 | 56 | 8.92E-14 | **** | [0.16, inf] | 1.28E+00 | 8.76E+10 | 1.00 |
| | TVAs | 0.16 ± 0.16 | 0.01 | 1.39E+01 | 181 | 8.43E-31 | **** | [0.14, inf] | 1.03E+00 | 3.76E+27 | 1.00 |
| s2 | LA1 | 0.04 ± 0.11 | 0.02 | 2.16E+00 | 32 | 1.94E-02 | * | [0.01, inf] | 3.80E-01 | 2.83E+00 | 0.68 |
| | RA1 | -0.01 ± 0.11 | 0.02 | n/a | n/a | n/a | n/a | n/a | n/a | n/a | n/a |
| | LmTVA | -0.02 ± 0.09 | 0.02 | n/a | n/a | n/a | n/a | n/a | n/a | n/a | n/a |
| | RmTVA | 0.03 ± 0.11 | 0.02 | 1.17E+00 | 21 | 1.27E-01 | ns | [-0.01, inf] | 2.50E-01 | 8.20E-01 | 0.31 |
| | LpTVA | -0.01 ± 0.10 | 0.02 | n/a | n/a | n/a | n/a | n/a | n/a | n/a | n/a |
| | RpTVA | 0.04 ± 0.10 | 0.03 | 1.38E+00 | 16 | 9.37E-02 | ns | [-0.01, inf] | 3.30E-01 | 1.11E+00 | 0.37 |
| | LaTVA | -0.05 ± 0.12 | 0.02 | n/a | n/a | n/a | n/a | n/a | n/a | n/a | n/a |
| | RaTVA | 0.03 ± 0.12 | 0.03 | 1.18E+00 | 19 | 1.26E-01 | ns | [-0.01, inf] | 2.60E-01 | 8.56E-01 | 0.31 |
| | A1 | 0.02 ± 0.11 | 0.01 | 1.19E+00 | 65 | 1.19E-01 | ns | [-0.01, inf] | 1.50E-01 | 5.31E-01 | 0.32 |
| | mTVA | 0.00 ± 0.10 | 0.02 | 5.00E-02 | 46 | 4.81E-01 | ns | [-0.02, inf] | 1.00E-02 | 3.17E-01 | 0.06 |
| | pTVA | 0.01 ± 0.10 | 0.02 | 5.10E-01 | 45 | 3.07E-01 | ns | [-0.02, inf] | 7.00E-02 | 3.61E-01 | 0.13 |
| | aTVA | -0.02 ± 0.12 | 0.02 | n/a | n/a | n/a | n/a | n/a | n/a | n/a | n/a |
| | TVAs | -0.00 ± 0.11 | 0.01 | n/a | n/a | n/a | n/a | n/a | n/a | n/a | n/a |
| s3 | LA1 | 0.04 ± 0.08 | 0.01 | 2.89E+00 | 32 | 3.39E-03 | ** | [0.02, inf] | 5.00E-01 | 1.21E+01 | 0.88 |
| | RA1 | 0.03 ± 0.13 | 0.02 | 1.48E+00 | 32 | 7.39E-02 | ns | [-0.00, inf] | 2.60E-01 | 1.01E+00 | 0.42 |
| | LmTVA | 0.04 ± 0.09 | 0.02 | 2.43E+00 | 28 | 1.10E-02 | * | [0.01, inf] | 4.50E-01 | 4.72E+00 | 0.76 |
| | RmTVA | 0.07 ± 0.09 | 0.02 | 4.38E+00 | 28 | 7.48E-05 | **** | [0.04, inf] | 8.10E-01 | 3.64E+02 | 1.00 |
| | LpTVA | 0.03 ± 0.12 | 0.02 | 1.48E+00 | 28 | 7.45E-02 | ns | [-0.00, inf] | 2.80E-01 | 1.06E+00 | 0.42 |
| | RpTVA | 0.04 ± 0.08 | 0.01 | 2.83E+00 | 35 | 3.87E-03 | ** | [0.02, inf] | 4.70E-01 | 1.05E+01 | 0.87 |
| | LaTVA | 0.09 ± 0.13 | 0.03 | 3.15E+00 | 23 | 2.24E-03 | ** | [0.04, inf] | 6.40E-01 | 1.91E+01 | 0.92 |
| | RaTVA | 0.07 ± 0.12 | 0.03 | 2.41E+00 | 17 | 1.38E-02 | * | [0.02, inf] | 5.70E-01 | 4.61E+00 | 0.75 |
| | A1 | 0.04 ± 0.11 | 0.01 | 2.80E+00 | 65 | 3.38E-03 | ** | [0.01, inf] | 3.40E-01 | 9.50E+00 | 0.87 |
| | mTVA | 0.06 ± 0.09 | 0.01 | 4.76E+00 | 57 | 6.83E-06 | **** | [0.04, inf] | 6.30E-01 | 2.76E+03 | 1.00 |
| | pTVA | 0.04 ± 0.10 | 0.01 | 2.92E+00 | 64 | 2.40E-03 | ** | [0.02, inf] | 3.60E-01 | 1.29E+01 | 0.89 |
| | aTVA | 0.08 ± 0.13 | 0.02 | 4.00E+00 | 41 | 1.28E-04 | *** | [0.05, inf] | 6.20E-01 | 2.05E+02 | 0.99 |
| | TVAs | 0.06 ± 0.11 | 0.01 | 6.62E+00 | 164 | 2.46E-10 | **** | [0.04, inf] | 5.20E-01 | 3.49E+07 | 1.00 |
| all | LA1 | 0.07 ± 0.12 | 0.01 | 5.58E+00 | 98 | 1.05E-07 | **** | [0.05, inf] | 5.60E-01 | 1.21E+05 | 1.00 |
| | RA1 | 0.08 ± 0.16 | 0.02 | 4.82E+00 | 98 | 2.60E-06 | **** | [0.05, inf] | 4.80E-01 | 5.85E+03 | 1.00 |
| | LmTVA | 0.13 ± 0.19 | 0.02 | 6.37E+00 | 86 | 4.45E-09 | **** | [0.10, inf] | 6.80E-01 | 2.55E+06 | 1.00 |
| | RmTVA | 0.09 ± 0.10 | 0.01 | 7.55E+00 | 77 | 3.72E-11 | **** | [0.07, inf] | 8.50E-01 | 2.55E+08 | 1.00 |
| | LpTVA | 0.03 ± 0.12 | 0.01 | 2.66E+00 | 90 | 4.59E-03 | ** | [0.01, inf] | 2.80E-01 | 6.39E+00 | 0.84 |
| | RpTVA | 0.04 ± 0.09 | 0.01 | 4.01E+00 | 84 | 6.63E-05 | **** | [0.02, inf] | 4.30E-01 | 3.00E+02 | 0.99 |
| | LaTVA | 0.11 ± 0.19 | 0.02 | 5.07E+00 | 83 | 1.20E-06 | **** | [0.07, inf] | 5.50E-01 | 1.27E+04 | 1.00 |
| | RaTVA | 0.07 ± 0.12 | 0.01 | 5.01E+00 | 63 | 2.34E-06 | **** | [0.05, inf] | 6.30E-01 | 7.30E+03 | 1.00 |
| | A1 | 0.07 ± 0.14 | 0.01 | 7.25E+00 | 197 | 4.67E-12 | **** | [0.06, inf] | 5.20E-01 | 1.54E+09 | 1.00 |
| | mTVA | 0.11 ± 0.16 | 0.01 | 9.12E+00 | 164 | 1.30E-16 | **** | [0.09, inf] | 7.10E-01 | 4.40E+13 | 1.00 |
| | pTVA | 0.04 ± 0.11 | 0.01 | 4.49E+00 | 175 | 6.53E-06 | **** | [0.02, inf] | 3.40E-01 | 2.00E+03 | 1.00 |
| | aTVA | 0.09 ± 0.17 | 0.01 | 6.81E+00 | 147 | 1.14E-10 | **** | [0.07, inf] | 5.60E-01 | 7.58E+07 | 1.00 |
| | TVAs | 0.08 ± 0.15 | 0.01 | 1.18E+01 | 488 | 9.58E-29 | **** | [0.07, inf] | 5.30E-01 | 2.93E+25 | 1.00 |

**Supplementary Table 2. Assessing the significance of brain encoding performance with LIN features.** This table reports the significance of the brain encoding performance with LIN features. We compared the distribution of Pearson's correlation coefficients to the chance level of 0.0 by conducting one-sample t-tests. Using a linear model, we calculated the correlation between the voxels in the speaker activity maps and the predicted voxels from the LIN features. s.e.m. = standard error of the mean. all = we combined the scores of all participants before computing the test. Here are reported the results of the statistical tests, t-value, degree of freedom (dof), p-value, degree of significance (unc. sig.), 95% confidence interval (CI95%), effect size (Cohen-d), Bayes Factor (BF10), and statistical power (power) for each participant and ROI.

| Subject | ROI | Correlation | s.e.m. | T | dof | p-val | unc. sig. | CI95% | cohen-d | BF10 | power |
|---|---|---|---|---|---|---|---|---|---|---|---|
| s1 | LA1 | 0.03 ± 0.11 | 0.02 | 1.46E+00 | 32 | 7.71E-02 | ns | [-0.00, inf] | 2.50E-01 | 9.75E-01 | 0.41 |
| | RA1 | 0.13 ± 0.09 | 0.02 | 8.06E+00 | 32 | 1.67E-09 | **** | [0.10, inf] | 1.40E+00 | 7.28E+06 | 1.00 |
| | LmTVA | 0.25 ± 0.16 | 0.03 | 8.95E+00 | 32 | 1.58E-10 | **** | [0.20, inf] | 1.56E+00 | 6.77E+07 | 1.00 |
| | RmTVA | 0.08 ± 0.09 | 0.02 | 4.89E+00 | 26 | 2.24E-05 | **** | [0.05, inf] | 9.40E-01 | 1.09E+03 | 1.00 |
| | LpTVA | -0.03 ± 0.12 | 0.02 | n/a | n/a | n/a | n/a | n/a | n/a | n/a | n/a |
| | RpTVA | -0.06 ± 0.11 | 0.02 | n/a | n/a | n/a | n/a | n/a | n/a | n/a | n/a |
| | LaTVA | 0.15 ± 0.16 | 0.03 | 5.34E+00 | 30 | 4.43E-06 | **** | [0.10, inf] | 9.60E-01 | 4.70E+03 | 1.00 |
| | RaTVA | 0.03 ± 0.11 | 0.02 | 1.55E+00 | 25 | 6.70E-02 | ns | [-0.00, inf] | 3.00E-01 | 1.19E+00 | 0.44 |
| | A1 | 0.08 ± 0.11 | 0.01 | 5.65E+00 | 65 | 1.93E-07 | **** | [0.06, inf] | 7.00E-01 | 7.57E+04 | 1.00 |
| | mTVA | 0.17 ± 0.15 | 0.02 | 8.69E+00 | 59 | 1.91E-12 | **** | [0.14, inf] | 1.12E+00 | 4.57E+09 | 1.00 |
| | pTVA | -0.04 ± 0.12 | 0.02 | n/a | n/a | n/a | n/a | n/a | n/a | n/a | n/a |
| | aTVA | 0.10 ± 0.15 | 0.02 | 4.94E+00 | 56 | 3.68E-06 | **** | [0.07, inf] | 6.50E-01 | 4.93E+03 | 1.00 |
| | TVAs | 0.07 ± 0.17 | 0.01 | 5.79E+00 | 181 | 1.52E-08 | **** | [0.05, inf] | 4.30E-01 | 6.37E+05 | 1.00 |
| s2 | LA1 | 0.04 ± 0.14 | 0.02 | 1.51E+00 | 32 | 7.01E-02 | ns | [-0.00, inf] | 2.60E-01 | 1.05E+00 | 0.43 |
| | RA1 | 0.01 ± 0.12 | 0.02 | 3.60E-01 | 32 | 3.59E-01 | ns | [-0.03, inf] | 6.00E-02 | 3.96E-01 | 0.10 |
| | LmTVA | 0.04 ± 0.07 | 0.01 | 3.07E+00 | 24 | 2.61E-03 | ** | [0.02, inf] | 6.10E-01 | 1.66E+01 | 0.91 |
| | RmTVA | 0.08 ± 0.10 | 0.02 | 3.95E+00 | 21 | 3.64E-04 | *** | [0.05, inf] | 8.40E-01 | 9.46E+01 | 0.99 |
| | LpTVA | -0.01 ± 0.10 | 0.02 | n/a | n/a | n/a | n/a | n/a | n/a | n/a | n/a |
| | RpTVA | 0.02 ± 0.13 | 0.03 | 7.30E-01 | 16 | 2.39E-01 | ns | [-0.03, inf] | 1.80E-01 | 6.29E-01 | 0.17 |
| | LaTVA | -0.01 ± 0.08 | 0.02 | n/a | n/a | n/a | n/a | n/a | n/a | n/a | n/a |
| | RaTVA | 0.02 ± 0.08 | 0.02 | 1.00E+00 | 19 | 1.64E-01 | ns | [-0.01, inf] | 2.20E-01 | 7.26E-01 | 0.25 |
| | A1 | 0.02 ± 0.13 | 0.02 | 1.38E+00 | 65 | 8.61E-02 | ns | [-0.00, inf] | 1.70E-01 | 6.66E-01 | 0.39 |
| | mTVA | 0.06 ± 0.09 | 0.01 | 4.92E+00 | 46 | 5.72E-06 | **** | [0.04, inf] | 7.20E-01 | 3.43E+03 | 1.00 |
| | pTVA | -0.00 ± 0.11 | 0.02 | n/a | n/a | n/a | n/a | n/a | n/a | n/a | n/a |
| | aTVA | 0.00 ± 0.08 | 0.01 | 4.10E-01 | 48 | 3.43E-01 | ns | [-0.02, inf] | 6.00E-02 | 3.36E-01 | 0.11 |
| | TVAs | 0.02 ± 0.10 | 0.01 | 2.65E+00 | 141 | 4.46E-03 | ** | [0.01, inf] | 2.20E-01 | 5.41E+00 | 0.84 |
| s3 | LA1 | 0.01 ± 0.09 | 0.02 | 3.50E-01 | 32 | 3.66E-01 | ns | [-0.02, inf] | 6.00E-02 | 3.94E-01 | 0.10 |
| | RA1 | 0.03 ± 0.11 | 0.02 | 1.62E+00 | 32 | 5.78E-02 | ns | [-0.00, inf] | 2.80E-01 | 1.21E+00 | 0.48 |
| | LmTVA | 0.05 ± 0.14 | 0.03 | 2.03E+00 | 28 | 2.61E-02 | * | [0.01, inf] | 3.80E-01 | 2.34E+00 | 0.63 |
| | RmTVA | 0.09 ± 0.08 | 0.02 | 5.64E+00 | 28 | 2.41E-06 | **** | [0.06, inf] | 1.05E+00 | 8.29E+03 | 1.00 |
| | LpTVA | 0.00 ± 0.10 | 0.02 | 2.20E-01 | 28 | 4.12E-01 | ns | [-0.03, inf] | 4.00E-02 | 4.04E-01 | 0.08 |
| | RpTVA | 0.01 ± 0.11 | 0.02 | 4.50E-01 | 35 | 3.30E-01 | ns | [-0.02, inf] | 7.00E-02 | 3.93E-01 | 0.11 |
| | LaTVA | 0.04 ± 0.12 | 0.03 | 1.60E+00 | 23 | 6.16E-02 | ns | [-0.00, inf] | 3.30E-01 | 1.31E+00 | 0.46 |
| | RaTVA | 0.11 ± 0.12 | 0.03 | 3.65E+00 | 17 | 9.96E-04 | *** | [0.06, inf] | 8.60E-01 | 4.13E+01 | 0.97 |
| | A1 | 0.02 ± 0.10 | 0.01 | 1.49E+00 | 65 | 7.09E-02 | ns | [-0.00, inf] | 1.80E-01 | 7.69E-01 | 0.43 |
| | mTVA | 0.07 ± 0.11 | 0.02 | 4.68E+00 | 57 | 9.11E-06 | **** | [0.05, inf] | 6.10E-01 | 2.12E+03 | 1.00 |
| | pTVA | 0.01 ± 0.11 | 0.01 | 4.90E-01 | 64 | 3.14E-01 | ns | [-0.02, inf] | 6.00E-02 | 3.05E-01 | 0.12 |
| | aTVA | 0.07 ± 0.13 | 0.02 | 3.53E+00 | 41 | 5.14E-04 | *** | [0.04, inf] | 5.50E-01 | 5.87E+01 | 0.97 |
| | TVAs | 0.05 ± 0.12 | 0.01 | 4.87E+00 | 164 | 1.32E-06 | **** | [0.03, inf] | 3.80E-01 | 9.31E+03 | 1.00 |
| all | LA1 | 0.02 ± 0.11 | 0.01 | 2.04E+00 | 98 | 2.19E-02 | * | [0.00, inf] | 2.10E-01 | 1.62E+00 | 0.65 |
| | RA1 | 0.06 ± 0.12 | 0.01 | 4.67E+00 | 98 | 4.87E-06 | **** | [0.04, inf] | 4.70E-01 | 3.24E+03 | 1.00 |
| | LmTVA | 0.12 ± 0.16 | 0.02 | 7.09E+00 | 86 | 1.77E-10 | **** | [0.09, inf] | 7.60E-01 | 5.59E+07 | 1.00 |
| | RmTVA | 0.09 ± 0.09 | 0.01 | 8.47E+00 | 77 | 6.43E-13 | **** | [0.07, inf] | 9.60E-01 | 1.27E+10 | 1.00 |
| | LpTVA | -0.01 ± 0.11 | 0.01 | n/a | n/a | n/a | n/a | n/a | n/a | n/a | n/a |
| | RpTVA | -0.02 ± 0.12 | 0.01 | n/a | n/a | n/a | n/a | n/a | n/a | n/a | n/a |
| | LaTVA | 0.07 ± 0.14 | 0.02 | 4.23E+00 | 83 | 2.96E-05 | **** | [0.04, inf] | 4.60E-01 | 6.36E+02 | 0.99 |
| | RaTVA | 0.05 ± 0.11 | 0.01 | 3.57E+00 | 63 | 3.50E-04 | *** | [0.03, inf] | 4.50E-01 | 7.23E+01 | 0.97 |
| | A1 | 0.04 ± 0.12 | 0.01 | 4.76E+00 | 197 | 1.89E-06 | **** | [0.03, inf] | 3.40E-01 | 6.19E+03 | 1.00 |
| | mTVA | 0.11 ± 0.13 | 0.01 | 1.01E+01 | 164 | 2.66E-19 | **** | [0.09, inf] | 7.90E-01 | 1.88E+16 | 1.00 |
| | pTVA | -0.01 ± 0.12 | 0.01 | n/a | n/a | n/a | n/a | n/a | n/a | n/a | n/a |
| | aTVA | 0.06 ± 0.13 | 0.01 | 5.52E+00 | 147 | 7.61E-08 | **** | [0.04, inf] | 4.50E-01 | 1.46E+05 | 1.00 |
| | TVAs | 0.05 ± 0.14 | 0.01 | 7.88E+00 | 488 | 1.05E-14 | **** | [0.04, inf] | 3.60E-01 | 4.33E+11 | 1.00 |

**Supplementary Table 3. Assessing the significance of brain encoding performance with VLS features.** This table reports the significance of the brain encoding performance with VLS features. We compared the distribution of Pearson's correlation coefficients to the chance level of 0.0 by conducting one-sample t-tests. Using a linear model, we calculated the correlation between the voxels in the speaker activity maps and the predicted voxels from the VLS features. s.e.m. = standard error of the mean. all = we combined the scores of all participants before computing the test. Here are reported the results of the statistical tests, t-value, degree of freedom (dof), p-value, degree of significance (unc. sig.), 95% confidence interval (CI95%), effect size (Cohen-d), Bayes Factor (BF10), and statistical power (power) for each participant and ROI.

| Subject | ROI | Correlation VLS | Correlation LIN | s.e.m. VLS | s.e.m. LIN | T VLS vs LIN | dof | p-val | unc. sig. | CI95% | cohen-d | BF10 | power |
|---|---|---|---|---|---|---|---|---|---|---|---|---|---|
| s1 | LA1 | 0.03 ± 0.11 | 0.13 ± 0.15 | 0.02 | 0.03 | -4.43E+00 | 32 | 1.03E-04 | *** | [-0.14, -0.05] | 7.30E-01 | 2.47E+02 | 0.98 |
| | RA1 | 0.13 ± 0.09 | 0.21 ± 0.14 | 0.02 | 0.03 | -3.75E+00 | 32 | 7.07E-04 | *** | [-0.11, -0.03] | 6.00E-01 | 4.39E+01 | 0.92 |
| | LmTVA | 0.25 ± 0.16 | 0.32 ± 0.13 | 0.03 | 0.02 | -3.90E+00 | 32 | 4.61E-04 | *** | [-0.11, -0.03] | 4.80E-01 | 6.43E+01 | 0.76 |
| | RmTVA | 0.08 ± 0.09 | 0.16 ± 0.07 | 0.02 | 0.01 | -5.48E+00 | 26 | 9.54E-06 | **** | [-0.10, -0.05] | 9.20E-01 | 2.24E+03 | 1.00 |
| | LpTVA | -0.03 ± 0.12 | 0.07 ± 0.13 | 0.02 | 0.02 | -6.49E+00 | 32 | 2.68E-07 | **** | [-0.13, -0.07] | 7.60E-01 | 5.95E+04 | 0.99 |
| | RpTVA | -0.06 ± 0.11 | 0.04 ± 0.08 | 0.02 | 0.02 | -5.09E+00 | 31 | 1.67E-05 | **** | [-0.14, -0.06] | 1.01E+00 | 1.31E+03 | 1.00 |
| | LaTVA | 0.15 ± 0.16 | 0.27 ± 0.15 | 0.03 | 0.03 | -7.34E+00 | 30 | 3.55E-08 | **** | [-0.15, -0.09] | 7.70E-01 | 3.95E+05 | 0.99 |
| | RaTVA | 0.03 ± 0.11 | 0.11 ± 0.10 | 0.02 | 0.02 | -4.24E+00 | 25 | 2.65E-04 | *** | [-0.11, -0.04] | 7.10E-01 | 1.11E+02 | 0.93 |
| | A1 | 0.08 ± 0.11 | 0.17 ± 0.15 | 0.01 | 0.02 | -5.81E+00 | 65 | 2.02E-07 | **** | [-0.12, -0.06] | 6.30E-01 | 6.96E+04 | 1.00 |
| | mTVA | 0.17 ± 0.15 | 0.25 ± 0.14 | 0.02 | 0.02 | -6.24E+00 | 59 | 5.16E-08 | **** | [-0.10, -0.05] | 5.00E-01 | 2.58E+05 | 0.97 |
| | pTVA | -0.04 ± 0.12 | 0.06 ± 0.11 | 0.02 | 0.01 | -8.06E+00 | 64 | 2.58E-11 | **** | [-0.12, -0.08] | 8.60E-01 | 3.62E+08 | 1.00 |
| | aTVA | 0.10 ± 0.15 | 0.20 ± 0.15 | 0.02 | 0.02 | -8.11E+00 | 56 | 5.09E-11 | **** | [-0.13, -0.08] | 6.60E-01 | 1.91E+08 | 1.00 |
| | TVAs | 0.07 ± 0.17 | 0.16 ± 0.16 | 0.01 | 0.01 | -1.29E+01 | 181 | 1.85E-27 | **** | [-0.11, -0.08] | 5.60E-01 | 1.89E+24 | 1.00 |
| s2 | LA1 | 0.04 ± 0.14 | 0.04 ± 0.11 | 0.02 | 0.02 | -2.70E-01 | 32 | 7.93E-01 | ns | [-0.03, 0.02] | 3.00E-02 | 1.92E-01 | 0.05 |
| | RA1 | 0.01 ± 0.12 | -0.01 ± 0.11 | 0.02 | 0.02 | 6.20E-01 | 32 | 5.38E-01 | ns | [-0.03, 0.06] | 1.30E-01 | 2.23E-01 | 0.11 |
| | LmTVA | 0.04 ± 0.07 | -0.02 ± 0.09 | 0.01 | 0.02 | 3.52E+00 | 24 | 1.77E-03 | ** | [0.03, 0.11] | 8.10E-01 | 2.11E+01 | 0.97 |
| | RmTVA | 0.08 ± 0.10 | 0.03 ± 0.11 | 0.02 | 0.02 | 3.74E+00 | 21 | 1.22E-03 | ** | [0.02, 0.09] | 5.20E-01 | 3.01E+01 | 0.65 |
| | LpTVA | -0.01 ± 0.10 | -0.01 ± 0.10 | 0.02 | 0.02 | -4.20E-01 | 28 | 6.78E-01 | ns | [-0.04, 0.02] | 6.00E-02 | 2.14E-01 | 0.06 |
| | RpTVA | 0.02 ± 0.13 | 0.04 ± 0.10 | 0.03 | 0.03 | -4.10E-01 | 16 | 6.88E-01 | ns | [-0.07, 0.05] | 1.00E-01 | 2.68E-01 | 0.07 |
| | LaTVA | -0.01 ± 0.08 | -0.05 ± 0.12 | 0.02 | 0.02 | 2.78E+00 | 28 | 9.51E-03 | ** | [0.01, 0.08] | 4.70E-01 | 4.75E+00 | 0.68 |
| | RaTVA | 0.02 ± 0.08 | 0.03 ± 0.12 | 0.02 | 0.03 | -4.30E-01 | 19 | 6.69E-01 | ns | [-0.07, 0.05] | 1.20E-01 | 2.53E-01 | 0.08 |
| | A1 | 0.02 ± 0.13 | 0.02 ± 0.11 | 0.02 | 0.01 | 4.00E-01 | 65 | 6.87E-01 | ns | [-0.02, 0.03] | 5.00E-02 | 1.46E-01 | 0.07 |
| | mTVA | 0.06 ± 0.09 | 0.00 ± 0.10 | 0.01 | 0.02 | 5.06E+00 | 46 | 7.24E-06 | **** | [0.04, 0.09] | 6.40E-01 | 2.62E+03 | 0.99 |
| | pTVA | -0.00 ± 0.11 | 0.01 ± 0.10 | 0.02 | 0.02 | -5.90E-01 | 45 | 5.57E-01 | ns | [-0.04, 0.02] | 8.00E-02 | 1.89E-01 | 0.08 |
| | aTVA | 0.00 ± 0.08 | -0.02 ± 0.12 | 0.01 | 0.02 | 1.50E+00 | 48 | 1.40E-01 | ns | [-0.01, 0.06] | 2.20E-01 | 4.43E-01 | 0.33 |
| | TVAs | 0.02 ± 0.10 | -0.00 ± 0.11 | 0.01 | 0.01 | 3.06E+00 | 141 | 2.64E-03 | ** | [0.01, 0.04] | 2.40E-01 | 8.00E+00 | 0.83 |
| s3 | LA1 | 0.01 ± 0.09 | 0.04 ± 0.08 | 0.02 | 0.01 | -2.32E+00 | 32 | 2.68E-02 | * | [-0.07, -0.00] | 4.20E-01 | 1.91E+00 | 0.64 |
| | RA1 | 0.03 ± 0.11 | 0.03 ± 0.13 | 0.02 | 0.02 | -1.00E-01 | 32 | 9.17E-01 | ns | [-0.04, 0.03] | 1.00E-02 | 1.87E-01 | 0.05 |
| | LmTVA | 0.05 ± 0.14 | 0.04 ± 0.09 | 0.03 | 0.02 | 7.20E-01 | 28 | 4.79E-01 | ns | [-0.02, 0.04] | 8.00E-02 | 2.50E-01 | 0.07 |
| | RmTVA | 0.09 ± 0.08 | 0.07 ± 0.09 | 0.02 | 0.02 | 9.30E-01 | 28 | 3.59E-01 | ns | [-0.02, 0.05] | 1.80E-01 | 2.94E-01 | 0.16 |
| | LpTVA | 0.00 ± 0.10 | 0.03 ± 0.12 | 0.02 | 0.02 | -1.82E+00 | 28 | 7.91E-02 | ns | [-0.06, 0.00] | 2.50E-01 | 8.47E-01 | 0.26 |
| | RpTVA | 0.01 ± 0.11 | 0.04 ± 0.08 | 0.02 | 0.01 | -2.26E+00 | 35 | 3.03E-02 | * | [-0.06, -0.00] | 3.10E-01 | 1.67E+00 | 0.44 |
| | LaTVA | 0.04 ± 0.12 | 0.09 ± 0.13 | 0.03 | 0.03 | -3.71E+00 | 23 | 1.15E-03 | ** | [-0.07, -0.02] | 3.70E-01 | 3.10E+01 | 0.40 |
| | RaTVA | 0.11 ± 0.12 | 0.07 ± 0.12 | 0.03 | 0.03 | 2.79E+00 | 17 | 1.25E-02 | * | [0.01, 0.07] | 3.00E-01 | 4.41E+00 | 0.23 |
| | A1 | 0.02 ± 0.10 | 0.04 ± 0.11 | 0.01 | 0.01 | -1.60E+00 | 65 | 1.14E-01 | ns | [-0.04, 0.00] | 1.80E-01 | 4.55E-01 | 0.29 |
| | mTVA | 0.07 ± 0.11 | 0.06 ± 0.09 | 0.02 | 0.01 | 1.19E+00 | 57 | 2.40E-01 | ns | [-0.01, 0.03] | 1.20E-01 | 2.79E-01 | 0.15 |
| | pTVA | 0.01 ± 0.11 | 0.04 ± 0.10 | 0.01 | 0.01 | -2.92E+00 | 64 | 4.88E-03 | ** | [-0.05, -0.01] | 2.80E-01 | 6.36E+00 | 0.61 |
| | aTVA | 0.07 ± 0.13 | 0.08 ± 0.13 | 0.02 | 0.02 | -9.50E-01 | 41 | 3.49E-01 | ns | [-0.03, 0.01] | 8.00E-02 | 2.54E-01 | 0.08 |
| | TVAs | 0.05 ± 0.12 | 0.06 ± 0.11 | 0.01 | 0.01 | -1.54E+00 | 164 | 1.25E-01 | ns | [-0.02, 0.00] | 9.00E-02 | 2.77E-01 | 0.20 |
| all | LA1 | 0.02 ± 0.11 | 0.07 ± 0.12 | 0.01 | 0.01 | -4.25E+00 | 98 | 4.92E-05 | **** | [-0.07, -0.02] | 3.80E-01 | 3.57E+02 | 0.97 |
| | RA1 | 0.06 ± 0.12 | 0.08 ± 0.16 | 0.01 | 0.02 | -1.64E+00 | 98 | 1.04E-01 | ns | [-0.04, 0.00] | 1.40E-01 | 4.06E-01 | 0.29 |
| | LmTVA | 0.12 ± 0.16 | 0.13 ± 0.19 | 0.02 | 0.02 | -4.10E-01 | 86 | 6.80E-01 | ns | [-0.03, 0.02] | 3.00E-02 | 1.29E-01 | 0.06 |
| | RmTVA | 0.09 ± 0.09 | 0.09 ± 0.10 | 0.01 | 0.01 | -4.00E-01 | 77 | 6.87E-01 | ns | [-0.03, 0.02] | 4.00E-02 | 1.35E-01 | 0.07 |
| | LpTVA | -0.01 ± 0.11 | 0.03 ± 0.12 | 0.01 | 0.01 | -4.81E+00 | 90 | 5.96E-06 | **** | [-0.07, -0.03] | 4.00E-01 | 2.64E+03 | 0.96 |
| | RpTVA | -0.02 ± 0.12 | 0.04 ± 0.09 | 0.01 | 0.01 | -4.60E+00 | 84 | 1.51E-05 | **** | [-0.08, -0.03] | 5.10E-01 | 1.13E+03 | 1.00 |
| | LaTVA | 0.07 ± 0.14 | 0.11 ± 0.19 | 0.02 | 0.02 | -3.42E+00 | 83 | 9.61E-04 | *** | [-0.07, -0.02] | 2.40E-01 | 2.46E+01 | 0.59 |
| | RaTVA | 0.05 ± 0.11 | 0.07 ± 0.12 | 0.01 | 0.01 | -1.81E+00 | 63 | 7.58E-02 | ns | [-0.05, 0.00] | 2.10E-01 | 6.31E-01 | 0.37 |
| | A1 | 0.04 ± 0.12 | 0.07 ± 0.14 | 0.01 | 0.01 | -4.02E+00 | 197 | 8.19E-05 | **** | [-0.05, -0.02] | 2.50E-01 | 1.70E+02 | 0.94 |
| | mTVA | 0.11 ± 0.13 | 0.11 ± 0.16 | 0.01 | 0.01 | -5.80E-01 | 164 | 5.64E-01 | ns | [-0.02, 0.01] | 3.00E-02 | 1.02E-01 | 0.07 |
| | pTVA | -0.01 ± 0.12 | 0.04 ± 0.11 | 0.01 | 0.01 | -6.65E+00 | 175 | 3.68E-10 | **** | [-0.06, -0.04] | 4.50E-01 | 2.27E+07 | 1.00 |
| | aTVA | 0.06 ± 0.13 | 0.09 ± 0.17 | 0.01 | 0.01 | -3.79E+00 | 147 | 2.23E-04 | *** | [-0.05, -0.02] | 2.30E-01 | 7.55E+01 | 0.78 |
| | TVAs | 0.05 ± 0.14 | 0.08 ± 0.15 | 0.01 | 0.01 | -6.28E+00 | 488 | 7.44E-10 | **** | [-0.04, -0.02] | 2.10E-01 | 7.90E+06 | 1.00 |

**Supplementary Table 4. Comparing the performance of brain encoding models.** This table reports the significance of the VLS-LIN difference in the brain encoding performance. We conducted paired t-tests between the brain encoding model's scores trained with the VLS features to predict the speaker activity maps' voxels and those trained with the LIN features. s.e.m. = standard error of the mean. all = we combined the scores of all participants before computing the test. Here are reported the results of the statistical tests, t-value, degree of freedom (dof), p-value, degree of significance (unc. sig.), 95% confidence interval (CI95%), effect size (Cohen-d), Bayes Factor (BF10), and statistical power (power) for each participant and ROI.

| Subject | Model | ROI | Correlation ROI | Correlation A1 | s.e.m. ROI | s.e.m. A1 | T ROI vs A1 | dof | p-val | unc. sig. | CI95% | cohen-d | BF10 | power |
|---|---|---|---|---|---|---|---|---|---|---|---|---|---|---|
| s1 | LIN | mTVA | 0.25 ± 0.14 | 0.17 ± 0.15 | 0.02 | 0.02 | 3.070000 | 124 | 2.62E-03 | ** | [0.03, 0.13] | 5.50E-01 | 1.25E+01 | 0.86 |
| | | pTVA | 0.06 ± 0.11 | 0.17 ± 0.15 | 0.01 | 0.02 | -4.710000 | 129 | 6.34E-06 | **** | [-0.16, -0.06] | 8.20E-01 | 2.60E+03 | 1.00 |
| | | aTVA | 0.20 ± 0.15 | 0.17 ± 0.15 | 0.02 | 0.02 | 1.150000 | 121 | 2.53E-01 | ns | [-0.02, 0.09] | 2.10E-01 | 3.50E-01 | 0.21 |
| | | TVAs | 0.16 ± 0.16 | 0.17 ± 0.15 | 0.01 | 0.02 | -0.130000 | 246 | 8.93E-01 | ns | [-0.05, 0.04] | 2.00E-02 | 1.57E-01 | 0.05 |
| | VLS | mTVA | 0.17 ± 0.15 | 0.08 ± 0.11 | 0.02 | 0.01 | 3.860000 | 124 | 1.81E-04 | *** | [0.05, 0.14] | 6.90E-01 | 1.28E+02 | 0.97 |
| | | pTVA | -0.04 ± 0.12 | 0.08 ± 0.11 | 0.02 | 0.01 | -6.020000 | 129 | 1.68E-08 | **** | [-0.17, -0.08] | 1.05E+00 | 6.23E+05 | 1.00 |
| | | aTVA | 0.10 ± 0.15 | 0.08 ± 0.11 | 0.02 | 0.01 | 0.750000 | 121 | 4.53E-01 | ns | [-0.03, 0.07] | 1.40E-01 | 2.49E-01 | 0.12 |
| | | TVAs | 0.07 ± 0.17 | 0.08 ± 0.11 | 0.01 | 0.01 | -0.350000 | 246 | 7.25E-01 | ns | [-0.05, 0.04] | 5.00E-02 | 1.65E-01 | 0.06 |
| s2 | LIN | mTVA | 0.00 ± 0.10 | 0.02 ± 0.11 | 0.02 | 0.01 | -0.760000 | 111 | 4.48E-01 | ns | [-0.06, 0.03] | 1.50E-01 | 2.62E-01 | 0.12 |
| | | pTVA | 0.01 ± 0.10 | 0.02 ± 0.11 | 0.02 | 0.01 | -0.430000 | 110 | 6.70E-01 | ns | [-0.05, 0.03] | 8.00E-02 | 2.21E-01 | 0.07 |
| | | aTVA | -0.02 ± 0.12 | 0.02 ± 0.11 | 0.02 | 0.01 | -1.580000 | 113 | 1.16E-01 | ns | [-0.08, 0.01] | 3.00E-01 | 6.15E-01 | 0.35 |
| | | TVAs | -0.00 ± 0.11 | 0.02 ± 0.11 | 0.01 | 0.01 | -1.220000 | 206 | 2.22E-01 | ns | [-0.05, 0.01] | 1.80E-01 | 3.24E-01 | 0.23 |
| | VLS | mTVA | 0.06 ± 0.09 | 0.02 ± 0.13 | 0.01 | 0.02 | 1.810000 | 111 | 7.29E-02 | ns | [-0.00, 0.08] | 3.50E-01 | 8.70E-01 | 0.43 |
| | | pTVA | -0.00 ± 0.11 | 0.02 ± 0.13 | 0.02 | 0.02 | -0.960000 | 110 | 3.41E-01 | ns | [-0.07, 0.02] | 1.80E-01 | 3.06E-01 | 0.16 |
| | | aTVA | 0.00 ± 0.08 | 0.02 ± 0.13 | 0.01 | 0.02 | -0.810000 | 113 | 4.20E-01 | ns | [-0.06, 0.03] | 1.50E-01 | 2.69E-01 | 0.13 |
| | | TVAs | 0.02 ± 0.10 | 0.02 ± 0.13 | 0.01 | 0.02 | -0.020000 | 206 | 9.87E-01 | ns | [-0.03, 0.03] | 0.00E+00 | 1.62E-01 | 0.05 |
| s3 | LIN | mTVA | 0.06 ± 0.09 | 0.04 ± 0.11 | 0.01 | 0.01 | 1.170000 | 122 | 2.43E-01 | ns | [-0.01, 0.06] | 2.10E-01 | 3.57E-01 | 0.21 |
| | | pTVA | 0.04 ± 0.10 | 0.04 ± 0.11 | 0.01 | 0.01 | -0.040000 | 129 | 9.71E-01 | ns | [-0.04, 0.03] | 1.00E-02 | 1.87E-01 | 0.05 |
| | | aTVA | 0.08 ± 0.13 | 0.04 ± 0.11 | 0.02 | 0.01 | 1.940000 | 106 | 5.55E-02 | ns | [-0.00, 0.09] | 3.80E-01 | 1.09E+00 | 0.48 |
| | | TVAs | 0.06 ± 0.11 | 0.04 ± 0.11 | 0.01 | 0.01 | 1.190000 | 229 | 2.35E-01 | ns | [-0.01, 0.05] | 1.70E-01 | 3.06E-01 | 0.22 |
| | VLS | mTVA | 0.07 ± 0.11 | 0.02 ± 0.10 | 0.02 | 0.01 | 2.700000 | 122 | 7.97E-03 | ** | [0.01, 0.09] | 4.90E-01 | 4.89E+00 | 0.76 |
| | | pTVA | 0.01 ± 0.11 | 0.02 ± 0.10 | 0.01 | 0.01 | -0.650000 | 129 | 5.16E-01 | ns | [-0.05, 0.02] | 1.10E-01 | 2.27E-01 | 0.10 |
| | | aTVA | 0.07 ± 0.13 | 0.02 ± 0.10 | 0.02 | 0.01 | 2.340000 | 106 | 2.11E-02 | * | [0.01, 0.10] | 4.60E-01 | 2.32E+00 | 0.64 |
| | | TVAs | 0.05 ± 0.12 | 0.02 ± 0.10 | 0.01 | 0.01 | 1.610000 | 229 | 1.08E-01 | ns | [-0.01, 0.06] | 2.30E-01 | 5.30E-01 | 0.36 |
| all | LIN | mTVA | 0.11 ± 0.16 | 0.07 ± 0.14 | 0.01 | 0.01 | 2.360000 | 361 | 1.86E-02 | * | [0.01, 0.07] | 2.50E-01 | 1.69E+00 | 0.65 |
| | | pTVA | 0.04 ± 0.11 | 0.07 ± 0.14 | 0.01 | 0.01 | -2.850000 | 372 | 4.57E-03 | ** | [-0.06, -0.01] | 3.00E-01 | 5.59E+00 | 0.81 |
| | | aTVA | 0.09 ± 0.17 | 0.07 ± 0.14 | 0.01 | 0.01 | 1.200000 | 344 | 2.29E-01 | ns | [-0.01, 0.05] | 1.30E-01 | 2.40E-01 | 0.22 |
| | | TVAs | 0.08 ± 0.15 | 0.07 ± 0.14 | 0.01 | 0.01 | 0.410000 | 685 | 6.79E-01 | ns | [-0.02, 0.03] | 3.00E-02 | 1.02E-01 | 0.07 |
| | VLS | mTVA | 0.11 ± 0.13 | 0.04 ± 0.12 | 0.01 | 0.01 | 4.910000 | 361 | 1.40E-06 | **** | [0.04, 0.09] | 5.20E-01 | 9.29E+03 | 1.00 |
| | | pTVA | -0.01 ± 0.12 | 0.04 ± 0.12 | 0.01 | 0.01 | -4.450000 | 372 | 1.13E-05 | **** | [-0.08, -0.03] | 4.60E-01 | 1.31E+03 | 0.99 |
| | | aTVA | 0.06 ± 0.13 | 0.04 ± 0.12 | 0.01 | 0.01 | 1.410000 | 344 | 1.58E-01 | ns | [-0.01, 0.05] | 1.50E-01 | 3.13E-01 | 0.29 |
| | | TVAs | 0.05 ± 0.14 | 0.04 ± 0.12 | 0.01 | 0.01 | 0.750000 | 685 | 4.56E-01 | ns | [-0.01, 0.03] | 6.00E-02 | 1.23E-01 | 0.12 |

**Supplementary Table 5. Comparing the performance of brain encoding ROIs.** This table reports the significance of the A1-TVAs difference in the brain encoding performance. We conducted two-sample t-tests between the brain encoding model's scores trained to predict A1 and those trained to predict temporal voice areas. s.e.m. = standard error of the mean. all = we combined the scores of all participants before computing the test. Here are reported the results of the statistical tests, t-value, degree of freedom (dof), p-value, degree of significance (unc. sig.), 95% confidence interval (CI95%), effect size (Cohen-d), Bayes Factor (BF10), and statistical power (power) for each participant and model.

| Subject | Model | ROI | Correlation | p-unc | p-corr | corr. sig. |
|---|---|---|---|---|---|---|
| s1 | LIN | LA1 | 0.07 | 1.39E-02 | 1.79E-01 | ns |
| | | RA1 | 0.08 | 4.20E-03 | 1.04E-01 | ns |
| | | LmTVA | 0.08 | 1.92E-02 | 3.80E-01 | ns |
| | | RmTVA | 0.06 | 7.14E-02 | 5.51E-01 | ns |
| | | LpTVA | 0.04 | 2.05E-01 | 6.53E-01 | ns |
| | | RpTVA | 0.03 | 3.16E-01 | 7.66E-01 | ns |
| | | LaTVA | 0.12 | 1.40E-03 | 5.04E-01 | ns |
| | | RaTVA | 0.07 | 4.26E-02 | 6.61E-01 | ns |
| | VLS | LA1 | 0.09 | 6.52E-02 | 6.53E-02 | ns |
| | | RA1 | 0.08 | 7.47E-02 | 7.49E-02 | ns |
| | | LmTVA | 0.11 | 1.28E-01 | 1.28E-01 | ns |
| | | RmTVA | 0.10 | 1.39E-01 | 1.39E-01 | ns |
| | | LpTVA | 0.09 | 1.94E-01 | 1.94E-01 | ns |
| | | RpTVA | 0.11 | 1.18E-01 | 1.18E-01 | ns |
| | | LaTVA | 0.19 | 4.17E-02 | 4.17E-02 | * |
| | | RaTVA | 0.13 | 1.30E-01 | 1.30E-01 | ns |
| s2 | LIN | LA1 | -0.01 | 5.03E-01 | 6.27E-01 | ns |
| | | RA1 | -0.00 | 1.87E-01 | 4.95E-01 | ns |
| | | LmTVA | 0.01 | 4.72E-01 | 7.19E-01 | ns |
| | | RmTVA | -0.01 | 7.98E-01 | 9.03E-01 | ns |
| | | LpTVA | -0.01 | 7.21E-01 | 8.13E-01 | ns |
| | | RpTVA | 0.00 | 4.07E-01 | 6.00E-01 | ns |
| | | LaTVA | -0.02 | 8.62E-01 | 9.22E-01 | ns |
| | | RaTVA | -0.02 | 7.69E-01 | 7.92E-01 | ns |
| | VLS | LA1 | 0.02 | 2.36E-01 | 2.52E-01 | ns |
| | | RA1 | 0.03 | 1.12E-01 | 1.12E-01 | ns |
| | | LmTVA | 0.06 | 2.29E-01 | 2.29E-01 | ns |
| | | RmTVA | -0.01 | 8.59E-01 | 9.26E-01 | ns |
| | | LpTVA | -0.02 | 8.89E-01 | 9.85E-01 | ns |
| | | RpTVA | 0.01 | 4.22E-01 | 4.54E-01 | ns |
| | | LaTVA | 0.03 | 3.37E-01 | 3.38E-01 | ns |
| | | RaTVA | 0.00 | 2.76E-01 | 3.23E-01 | ns |
| s3 | LIN | LA1 | -0.00 | 5.71E-01 | 6.66E-01 | ns |
| | | RA1 | 0.05 | 3.00E-04 | 5.00E-02 | * |
| | | LmTVA | 0.05 | 4.10E-03 | 2.04E-01 | ns |
| | | RmTVA | 0.05 | 2.20E-03 | 1.16E-01 | ns |
| | | LpTVA | 0.05 | 5.80E-03 | 1.73E-01 | ns |
| | | RpTVA | 0.04 | 2.66E-02 | 4.60E-01 | ns |
| | | LaTVA | 0.12 | 0.00E+00 | 7.70E-02 | ns |
| | | RaTVA | 0.03 | 3.35E-02 | 3.26E-01 | ns |
| | VLS | LA1 | 0.02 | 1.78E-01 | 2.10E-01 | ns |
| | | RA1 | 0.07 | 1.42E-02 | 1.42E-02 | * |
| | | LmTVA | 0.11 | 7.20E-03 | 7.20E-03 | ** |
| | | RmTVA | 0.05 | 1.23E-01 | 1.23E-01 | ns |
| | | LpTVA | 0.08 | 5.82E-02 | 5.82E-02 | ns |
| | | RpTVA | 0.13 | 1.56E-02 | 1.56E-02 | * |
| | | LaTVA | 0.23 | 1.00E-04 | 1.00E-04 | **** |
| | | RaTVA | 0.04 | 2.34E-01 | 2.34E-01 | ns |

**Supplementary Table 6. Assessing the significance of the RSA brain-model correlation.** This table reports the significance of the RSA brain-model performance. The brain-model correlation coefficients were computed between the ranked representational dissimilarity matrices. The correlation was compared to 0 using a 'maximum statistics' approach in which they are compared to a distribution of correlation coefficients drawn from a large number of random permutations of the model RDMs' rows and columns while controlling for the number of comparisons performed (cf. Methods) (Maris & Oostenveld, 2007), for each participant, model and ROI.

| Subject | ROI | Correlation VLS | Correlation LIN | p-corr | p-unc | corr. sig. |
|---|---|---|---|---|---|---|
| s1 | LA1 | 0.09 | 0.07 | 4.45E-01 | 2.99E-01 | ns |
| | RA1 | 0.08 | 0.08 | 8.30E-01 | 5.05E-01 | ns |
| | LmTVA | 0.11 | 0.08 | 4.63E-01 | 4.51E-01 | ns |
| | RmTVA | 0.10 | 0.06 | 3.98E-01 | 3.97E-01 | ns |
| | LpTVA | 0.09 | 0.04 | 2.86E-01 | 2.84E-01 | ns |
| | RpTVA | 0.11 | 0.03 | 1.11E-01 | 1.11E-01 | ns |
| | LaTVA | 0.19 | 0.12 | 3.94E-01 | 3.94E-01 | ns |
| | RaTVA | 0.13 | 0.07 | 3.48E-01 | 3.48E-01 | ns |
| s2 | LA1 | 0.02 | -0.01 | 3.25E-01 | 1.65E-01 | ns |
| | RA1 | 0.03 | -0.00 | 1.58E-01 | 1.41E-01 | ns |
| | LmTVA | 0.06 | 0.01 | 1.78E-01 | 1.72E-01 | ns |
| | RmTVA | -0.01 | -0.01 | 1.00E+00 | 8.15E-01 | ns |
| | LpTVA | -0.02 | -0.01 | 1.00E+00 | 8.72E-01 | ns |
| | RpTVA | 0.01 | 0.00 | 7.13E-01 | 4.47E-01 | ns |
| | LaTVA | 0.03 | -0.02 | 1.20E-01 | 1.19E-01 | ns |
| | RaTVA | 0.00 | -0.02 | 3.94E-01 | 1.13E-01 | ns |
| s3 | LA1 | 0.02 | -0.00 | 3.22E-01 | 1.05E-01 | ns |
| | RA1 | 0.07 | 0.05 | 4.83E-01 | 3.22E-01 | ns |
| | LmTVA | 0.11 | 0.05 | 6.61E-02 | 6.25E-02 | ns |
| | RmTVA | 0.05 | 0.05 | 1.00E+00 | 5.38E-01 | ns |
| | LpTVA | 0.08 | 0.05 | 4.30E-01 | 3.08E-01 | ns |
| | RpTVA | 0.13 | 0.04 | 3.66E-02 | 3.66E-02 | * |
| | LaTVA | 0.23 | 0.12 | 1.75E-02 | 1.75E-02 | * |
| | RaTVA | 0.04 | 0.03 | 7.67E-01 | 6.08E-01 | ns |

**Supplementary Table 7. Comparing the performance of the RSA models.** This table reports the significance of the RSA brain-model difference. We compared the correlation coefficients between brain RDM and VLS RDM with those from the brain RDM and LIN RDM within participants and hemispheres using one-tailed tests, based on the a priori hypothesis that the VLS models would exhibit greater brain-model correlations than the LIN models (cf. Methods).

| Model | ROI | Accuracy (%) | s.e.m. | T | dof | p-val | unc. sig. | CI95% | cohen-d | BF10 | power |
|---|---|---|---|---|---|---|---|---|---|---|---|
| LIN | LA1 | 43.33 ± 2.22 | 0.51 | n/a | n/a | n/a | n/a | n/a | n/a | n/a | n/a |
| | RA1 | 50.83 ± 1.98 | 0.46 | 1.83E+00 | 19 | 4.14E-02 | * | [50.05, inf] | 4.10E-01 | 1.89E+00 | 0.55 |
| | LmTVA | 38.89 ± 0.00 | 0.00 | n/a | n/a | n/a | n/a | n/a | n/a | n/a | n/a |
| | RmTVA | 61.39 ± 1.21 | 0.28 | 4.10E+01 | 19 | 2.61E-20 | **** | [60.91, inf] | 9.17E+00 | 1.04E+17 | 1.00 |
| | LpTVA | 66.67 ± 0.02 | 0.00 | 4.59E+03 | 19 | 3.32E-59 | **** | [66.66, inf] | 1.03E+03 | 6.88E+33 | 1.00 |
| | RpTVA | 77.50 ± 1.21 | 0.28 | 9.90E+01 | 19 | 1.51E-27 | **** | [77.02, inf] | 2.21E+01 | 7.38E+23 | 1.00 |
| | LaTVA | 44.44 ± 0.00 | 0.00 | n/a | n/a | n/a | n/a | n/a | n/a | n/a | n/a |
| | RaTVA | 44.44 ± 0.00 | 0.00 | n/a | n/a | n/a | n/a | n/a | n/a | n/a | n/a |
| | A1 | 47.08 ± 4.30 | 0.69 | n/a | n/a | n/a | n/a | n/a | n/a | n/a | n/a |
| | mTVA | 50.14 ± 11.28 | 1.81 | 8.00E-02 | 39 | 4.70E-01 | ns | [47.09, inf] | 1.00E-02 | 3.42E-01 | 0.06 |
| | pTVA | 72.08 ± 5.48 | 0.88 | 2.51E+01 | 39 | 5.18E-26 | **** | [70.60, inf] | 3.98E+00 | 5.89E+22 | 1.00 |
| | aTVA | 44.44 ± 0.00 | 0.00 | n/a | n/a | n/a | n/a | n/a | n/a | n/a | n/a |
| | TVAs | 55.56 ± 13.94 | 1.28 | 4.35E+00 | 119 | 1.47E-05 | **** | [53.44, inf] | 4.00E-01 | 1.08E+03 | 1.00 |
| VLS | LA1 | 61.94 ± 1.98 | 0.46 | 2.62E+01 | 19 | 1.08E-16 | **** | [61.16, inf] | 5.87E+00 | 3.93E+13 | 1.00 |
| | RA1 | 60.28 ± 1.98 | 0.46 | 2.26E+01 | 19 | 1.73E-15 | **** | [59.49, inf] | 5.05E+00 | 2.88E+12 | 1.00 |
| | LmTVA | 55.56 ± 0.02 | 0.00 | 1.53E+03 | 19 | 3.86E-50 | **** | [55.55, inf] | 3.42E+02 | 4.95E+33 | 1.00 |
| | RmTVA | 44.44 ± 0.00 | 0.00 | n/a | n/a | n/a | n/a | n/a | n/a | n/a | n/a |
| | LpTVA | 66.67 ± 0.02 | 0.00 | 4.59E+03 | 19 | 3.32E-59 | **** | [66.66, inf] | 1.03E+03 | 6.88E+33 | 1.00 |
| | RpTVA | 61.11 ± 0.02 | 0.00 | 3.06E+03 | 19 | 7.36E-56 | **** | [61.10, inf] | 6.85E+02 | 6.53E+33 | 1.00 |
| | LaTVA | 50.83 ± 1.98 | 0.46 | 1.83E+00 | 19 | 4.14E-02 | * | [50.05, inf] | 4.10E-01 | 1.89E+00 | 0.55 |
| | RaTVA | 44.17 ± 1.21 | 0.28 | n/a | n/a | n/a | n/a | n/a | n/a | n/a | n/a |
| | A1 | 61.11 ± 2.15 | 0.34 | 3.22E+01 | 39 | 4.92E-30 | **** | [60.53, inf] | 5.10E+00 | 4.78E+26 | 1.00 |
| | mTVA | 50.00 ± 5.56 | 0.89 | n/a | n/a | n/a | n/a | n/a | n/a | n/a | n/a |
| | pTVA | 63.89 ± 2.78 | 0.44 | 3.12E+01 | 39 | 1.65E-29 | **** | [63.14, inf] | 4.94E+00 | 1.47E+26 | 1.00 |
| | aTVA | 47.50 ± 3.72 | 0.60 | n/a | n/a | n/a | n/a | n/a | n/a | n/a | n/a |
| | TVAs | 53.80 ± 8.33 | 0.76 | 4.97E+00 | 119 | 1.14E-06 | **** | [52.53, inf] | 4.50E-01 | 1.20E+04 | 1.00 |

**Supplementary Table 8. Assessing the significance of speaker gender decoding performance using VLS and LIN models based on voxel activity.** This table reports the significance of the speaker's gender decoding performance. Linear classifiers were pre-trained to detect speaker gender (2 classes) from either the VLS or the LIN models. The speaker gender of the 18 Test Stimuli (3 participants x 6 stimuli per participant) was classified using either the VLS coordinates, or the LIN features with these classifiers. We used one-sample t-tests to compare the mean of the accuracy distribution across 20 random classifier initializations (20 classifiers trained with a different initialization seed) with a chance level of 50%. s.e.m. = standard error of the mean. Here are reported the results of the statistical tests, t-value, degree of freedom (dof), p-value, degree of significance (unc. sig.), 95% confidence interval (CI95%), effect size (Cohen-d), Bayes Factor (BF10), and statistical power (power) for each model and ROI.

| Model | ROI | Accuracy (%) | s.e.m. | T | dof | p-val | unc. sig. | CI95% | cohen-d | BF10 | power |
|---|---|---|---|---|---|---|---|---|---|---|---|
| LIN | LA1 | 50.42 ± 4.15 | 0.95 | 4.40E-01 | 19 | 3.33E-01 | ns | [48.77, inf] | 1.00E-01 | 5.07E-01 | 0.11 |
| | RA1 | 10.83 ± 3.82 | 0.88 | n/a | n/a | n/a | n/a | n/a | n/a | n/a | n/a |
| | LmTVA | 44.17 ± 3.82 | 0.88 | n/a | n/a | n/a | n/a | n/a | n/a | n/a | n/a |
| | RmTVA | 50.42 ± 6.71 | 1.54 | 2.70E-01 | 19 | 3.95E-01 | ns | [47.76, inf] | 6.00E-02 | 4.80E-01 | 0.08 |
| | LpTVA | 52.50 ± 3.82 | 0.88 | 2.85E+00 | 19 | 5.08E-03 | ** | [50.99, inf] | 6.40E-01 | 1.01E+01 | 0.87 |
| | RpTVA | 56.67 ± 3.33 | 0.76 | 8.72E+00 | 19 | 2.29E-08 | **** | [55.34, inf] | 1.95E+00 | 6.13E+05 | 1.00 |
| | LaTVA | 52.50 ± 3.82 | 0.88 | 2.85E+00 | 19 | 5.08E-03 | ** | [50.99, inf] | 6.40E-01 | 1.01E+01 | 0.87 |
| | RaTVA | 75.42 ± 6.17 | 1.41 | 1.80E+01 | 19 | 1.11E-13 | **** | [72.97, inf] | 4.02E+00 | 5.71E+10 | 1.00 |
| | A1 | 30.62 ± 20.19 | 3.23 | n/a | n/a | n/a | n/a | n/a | n/a | n/a | n/a |
| | mTVA | 47.29 ± 6.29 | 1.01 | n/a | n/a | n/a | n/a | n/a | n/a | n/a | n/a |
| | pTVA | 54.58 ± 4.15 | 0.66 | 6.90E+00 | 39 | 1.45E-08 | **** | [53.46, inf] | 1.09E+00 | 9.41E+05 | 1.00 |
| | aTVA | 63.96 ± 12.55 | 2.01 | 6.94E+00 | 39 | 1.28E-08 | **** | [60.57, inf] | 1.10E+00 | 1.06E+06 | 1.00 |
| | TVAs | 55.28 ± 10.86 | 1.00 | 5.30E+00 | 119 | 2.69E-07 | **** | [53.63, inf] | 4.80E-01 | 4.69E+04 | 1.00 |
| VLS | LA1 | 66.67 ± 0.02 | 0.00 | 4.59E+03 | 19 | 3.32E-59 | **** | [66.66, inf] | 1.03E+03 | 6.88E+33 | 1.00 |
| | RA1 | 8.33 ± 0.00 | 0.00 | n/a | n/a | n/a | n/a | n/a | n/a | n/a | n/a |
| | LmTVA | 49.17 ± 12.61 | 2.89 | n/a | n/a | n/a | n/a | n/a | n/a | n/a | n/a |
| | RmTVA | 41.67 ± 0.00 | 0.00 | n/a | n/a | n/a | n/a | n/a | n/a | n/a | n/a |
| | LpTVA | 58.33 ± 0.02 | 0.00 | 2.30E+03 | 19 | 1.74E-53 | **** | [58.33, inf] | 5.14E+02 | 6.07E+33 | 1.00 |
| | RpTVA | 71.67 ± 4.08 | 0.94 | 2.31E+01 | 19 | 1.11E-15 | **** | [70.05, inf] | 5.17E+00 | 4.36E+12 | 1.00 |
| | LaTVA | 56.67 ± 3.33 | 0.76 | 8.72E+00 | 19 | 2.29E-08 | **** | [55.34, inf] | 1.95E+00 | 6.13E+05 | 1.00 |
| | RaTVA | 64.17 ± 3.82 | 0.88 | 1.62E+01 | 19 | 7.29E-13 | **** | [62.65, inf] | 3.62E+00 | 9.69E+09 | 1.00 |
| | A1 | 37.50 ± 29.17 | 4.67 | n/a | n/a | n/a | n/a | n/a | n/a | n/a | n/a |
| | mTVA | 45.42 ± 9.67 | 1.55 | n/a | n/a | n/a | n/a | n/a | n/a | n/a | n/a |
| | pTVA | 65.00 ± 7.26 | 1.16 | 1.29E+01 | 39 | 6.05E-16 | **** | [63.04, inf] | 2.04E+00 | 1.05E+13 | 1.00 |
| | aTVA | 60.42 ± 5.19 | 0.83 | 1.25E+01 | 39 | 1.46E-15 | **** | [59.02, inf] | 1.98E+00 | 4.51E+12 | 1.00 |
| | TVAs | 56.94 ± 11.30 | 1.04 | 6.70E+00 | 119 | 3.59E-10 | **** | [55.23, inf] | 6.10E-01 | 2.64E+07 | 1.00 |

**Supplementary Table 9. Assessing the significance of speaker age decoding performance using VLS and LIN models based on voxel activity.** This table reports the significance of the speaker age decoding performance. Linear classifiers were pre-trained to detect speaker age (2 classes) from either the VLS or the LIN models. The speaker age of the 18 Test Stimuli (3 participants x 6 stimuli per participant) was classified using either the VLS or LIN coordinates with these classifiers. We used one-sample t-tests to compare the mean of the accuracy distribution across 20 random classifier initializations (20 classifiers trained with a different initialization seed) with the chance level of 50%. s.e.m. = standard error of the mean. Here are reported the results of the statistical tests, t-value, degree of freedom (dof), p-value, and degree of significance (unc. sig.), 95% confidence interval (CI95%), effect size (Cohen-d), Bayes Factor (BF10), and statistical power (power) for each model and ROI.

| Model | ROI | Accuracy (%) | s.e.m. | T | dof | p-val | unc. sig. | CI95% | cohen-d | BF10 | power |
|---|---|---|---|---|---|---|---|---|---|---|---|
| LIN | LA1 | 0.29 ± 1.28 | 0.29 | n/a | n/a | n/a | n/a | n/a | n/a | n/a | n/a |
| | RA1 | 18.09 ± 3.26 | 0.75 | 1.63E+01 | 19 | 6.14E-13 | **** | [16.80, inf] | 3.65E+00 | 1.14E+10 | 1.00 |
| | LmTVA | 11.18 ± 4.01 | 0.92 | 5.75E+00 | 19 | 7.61E-06 | **** | [9.59, inf] | 1.29E+00 | 3.01E+03 | 1.00 |
| | RmTVA | 2.35 ± 3.03 | 0.69 | n/a | n/a | n/a | n/a | n/a | n/a | n/a | n/a |
| | LpTVA | 12.21 ± 3.39 | 0.78 | 8.13E+00 | 19 | 6.54E-08 | **** | [10.86, inf] | 1.82E+00 | 2.32E+05 | 1.00 |
| | RpTVA | 6.76 ± 4.66 | 1.07 | 8.30E-01 | 19 | 2.10E-01 | ns | [4.92, inf] | 1.80E-01 | 6.29E-01 | 0.20 |
| | LaTVA | 11.47 ± 1.28 | 0.29 | 1.90E+01 | 19 | 4.04E-14 | **** | [10.96, inf] | 4.25E+00 | 1.48E+11 | 1.00 |
| | RaTVA | 7.35 ± 8.29 | 1.90 | 7.70E-01 | 19 | 2.25E-01 | ns | [4.06, inf] | 1.70E-01 | 6.07E-01 | 0.18 |
| | A1 | 9.19 ± 9.24 | 1.48 | 2.24E+00 | 39 | 1.55E-02 | * | [6.70, inf] | 3.50E-01 | 3.15E+00 | 0.71 |
| | mTVA | 6.76 ± 5.67 | 0.91 | 9.70E-01 | 39 | 1.68E-01 | ns | [5.24, inf] | 1.50E-01 | 5.30E-01 | 0.25 |
| | pTVA | 9.49 ± 4.90 | 0.78 | 4.59E+00 | 39 | 2.24E-05 | **** | [8.16, inf] | 7.30E-01 | 1.01E+03 | 1.00 |
| | aTVA | 9.41 ± 6.28 | 1.01 | 3.51E+00 | 39 | 5.75E-04 | *** | [7.72, inf] | 5.50E-01 | 5.39E+01 | 0.96 |
| | TVAs | 8.55 ± 5.78 | 0.53 | 5.04E+00 | 119 | 8.44E-07 | **** | [7.68, inf] | 4.60E-01 | 1.59E+04 | 1.00 |
| VLS | LA1 | 0.15 ± 0.64 | 0.15 | n/a | n/a | n/a | n/a | n/a | n/a | n/a | n/a |
| | RA1 | 11.47 ± 5.09 | 1.17 | 4.79E+00 | 19 | 6.37E-05 | **** | [9.45, inf] | 1.07E+00 | 4.49E+02 | 1.00 |
| | LmTVA | 11.47 ± 4.73 | 1.09 | 5.15E+00 | 19 | 2.87E-05 | **** | [9.59, inf] | 1.15E+00 | 9.13E+02 | 1.00 |
| | RmTVA | 0.59 ± 1.50 | 0.34 | n/a | n/a | n/a | n/a | n/a | n/a | n/a | n/a |
| | LpTVA | 9.71 ± 3.37 | 0.77 | 4.95E+00 | 19 | 4.44E-05 | **** | [8.37, inf] | 1.11E+00 | 6.19E+02 | 1.00 |
| | RpTVA | 22.65 ± 2.10 | 0.48 | 3.48E+01 | 19 | 5.69E-19 | **** | [21.81, inf] | 7.78E+00 | 5.62E+15 | 1.00 |
| | LaTVA | 10.29 ± 4.51 | 1.03 | 4.27E+00 | 19 | 2.09E-04 | *** | [8.51, inf] | 9.50E-01 | 1.57E+02 | 0.99 |
| | RaTVA | 6.18 ± 3.94 | 0.90 | 3.30E-01 | 19 | 3.74E-01 | ns | [4.62, inf] | 7.00E-02 | 4.88E-01 | 0.09 |
| | A1 | 5.81 ± 6.72 | 1.08 | n/a | n/a | n/a | n/a | n/a | n/a | n/a | n/a |
| | mTVA | 6.03 ± 6.48 | 1.04 | 1.40E-01 | 39 | 4.44E-01 | ns | [4.28, inf] | 2.00E-02 | 3.44E-01 | 0.07 |
| | pTVA | 16.18 ± 7.05 | 1.13 | 9.12E+00 | 39 | 1.65E-11 | **** | [14.27, inf] | 1.44E+00 | 5.85E+08 | 1.00 |
| | aTVA | 8.24 ± 4.71 | 0.75 | 3.12E+00 | 39 | 1.69E-03 | ** | [6.97, inf] | 4.90E-01 | 2.09E+01 | 0.92 |
| | TVAs | 10.15 ± 7.55 | 0.69 | 6.17E+00 | 119 | 4.96E-09 | **** | [9.00, inf] | 5.60E-01 | 2.12E+06 | 1.00 |

**Supplementary Table 10. Assessing the significance of speaker identity decoding performance using VLS and LIN models based on voxel activity.** This table reports the significance of the speaker speaker decoding performance. Linear classifiers were pre-trained to detect speaker identity (17 classes) from either the VLS or the LIN models. The speaker identity of the 18 Test Stimuli (3 participants x 6 stimuli per participant) was classified using either the VLS or LIN coordinates with these classifiers. We used one-sample t-tests to compare the mean of the accuracy distribution across 20 random classifier initializations (20 classifiers trained with a different initialization seed) with the chance level of 5.88%. s.e.m. = standard error of the mean. Here are reported the results of the statistical tests, t-value, degree of freedom (dof), p-value, and degree of significance (unc. sig.), 95% confidence interval (CI95%), effect size (Cohen-d), Bayes Factor (BF10), and statistical power (power) for each model and ROI.

| Category | ROI | Accuracy VLS (%) | Accuracy LIN (%) | s.e.m. VLS | s.e.m. LIN | T VLS vs LIN | dof | p-val | unc. sig. | CI95% | cohen-d | BF10 | power |
|---|---|---|---|---|---|---|---|---|---|---|---|---|---|
| Gender | LA1 | 61.94 ± 1.98 | 43.33 ± 2.22 | 0.46 | 0.51 | 3.06E+01 | 19 | 1.24E-17 | **** | [17.34, 19.88] | 8.610000 | 2.94E+14 | 1.00 |
| | RA1 | 60.28 ± 1.98 | 50.83 ± 1.98 | 0.46 | 0.46 | 1.62E+01 | 19 | 1.46E-12 | **** | [8.22, 10.67] | 4.640000 | 4.85E+09 | 1.00 |
| | LmTVA | 55.56 ± 0.02 | 38.89 ± 0.02 | 0.00 | 0.00 | INF | 19 | 0.00E+00 | **** | [nan, nan] | 1027.400000 | nan | 1.00 |
| | RmTVA | 44.44 ± 0.02 | 61.39 ± 1.21 | 0.00 | 0.28 | -6.10E+01 | 19 | 2.91E-23 | **** | [-17.53, -16.36] | 19.290000 | 6.23E+19 | 1.00 |
| | LpTVA | 66.67 ± 0.02 | 66.67 ± 0.02 | 0.00 | 0.00 | nan | 19 | nan | ns | [nan, nan] | 0.000000 | nan | 0.05 |
| | RpTVA | 61.11 ± 0.02 | 77.50 ± 1.21 | 0.00 | 0.28 | -5.90E+01 | 19 | 5.47E-23 | **** | [-16.97, -15.81] | 18.660000 | 3.43E+19 | 1.00 |
| | LaTVA | 50.83 ± 1.98 | 44.44 ± 0.02 | 0.46 | 0.00 | 1.41E+01 | 19 | 1.65E-11 | **** | [5.44, 7.34] | 4.440000 | 4.96E+08 | 1.00 |
| | RaTVA | 44.17 ± 1.21 | 44.44 ± 0.02 | 0.28 | 0.00 | -1.00E+00 | 19 | 3.30E-01 | ns | [-0.86, 0.30] | 0.320000 | 3.61E-01 | 0.27 |
| | A1 | 61.11 ± 2.15 | 47.08 ± 4.30 | 0.34 | 0.69 | 1.66E+01 | 39 | 2.67E-19 | **** | [12.32, 15.73] | 4.070000 | 1.78E+16 | 1.00 |
| | mTVA | 50.00 ± 5.56 | 50.14 ± 11.28 | 0.89 | 1.81 | -5.00E-02 | 39 | 9.59E-01 | ns | [-5.59, 5.31] | 0.020000 | 1.71E-01 | 0.05 |
| | pTVA | 63.89 ± 2.78 | 72.08 ± 5.48 | 0.44 | 0.88 | -6.21E+00 | 39 | 2.64E-07 | **** | [-10.86, -5.53] | 1.860000 | 5.92E+04 | 1.00 |
| | aTVA | 47.50 ± 3.72 | 44.44 ± 0.01 | 0.60 | 0.00 | 5.14E+00 | 39 | 8.05E-06 | **** | [1.85, 4.26] | 1.150000 | 2.45E+03 | 1.00 |
| | TVAs | 53.80 ± 8.33 | 55.56 ± 13.94 | 0.76 | 1.28 | -1.60E+00 | 119 | 1.12E-01 | ns | [-3.94, 0.42] | 0.150000 | 3.49E-01 | 0.38 |
| Age | LA1 | 66.67 ± 0.02 | 50.42 ± 4.15 | 0.00 | 0.95 | 1.71E+01 | 19 | 5.58E-13 | **** | [14.26, 18.24] | 5.400000 | 1.20E+10 | 1.00 |
| | RA1 | 8.33 ± 0.02 | 10.83 ± 3.82 | 0.00 | 0.88 | -2.85E+00 | 19 | 1.03E-02 | * | [-4.34, -0.66] | 0.900000 | 5.02E+00 | 0.97 |
| | LmTVA | 49.17 ± 12.61 | 44.17 ± 3.82 | 2.89 | 0.88 | 1.71E+00 | 19 | 1.04E-01 | ns | [-1.12, 11.12] | 0.520000 | 7.97E-01 | 0.60 |
| | RmTVA | 41.67 ± 0.02 | 50.42 ± 6.71 | 0.00 | 1.54 | -5.69E+00 | 19 | 1.76E-05 | **** | [-11.97, -5.53] | 1.800000 | 1.32E+03 | 1.00 |
| | LpTVA | 58.33 ± 0.02 | 52.50 ± 3.82 | 0.00 | 0.88 | 6.67E+00 | 19 | 2.24E-06 | **** | [4.00, 7.66] | 2.110000 | 8.58E+03 | 1.00 |
| | RpTVA | 71.67 ± 4.08 | 56.67 ± 3.33 | 0.94 | 0.76 | 1.16E+01 | 19 | 4.80E-10 | **** | [12.29, 17.71] | 3.920000 | 2.12E+07 | 1.00 |
| | LaTVA | 56.67 ± 3.33 | 52.50 ± 3.82 | 0.76 | 0.88 | 3.68E+00 | 19 | 1.58E-03 | ** | [1.80, 6.53] | 1.130000 | 2.46E+01 | 1.00 |
| | RaTVA | 64.17 ± 3.82 | 75.42 ± 6.17 | 0.88 | 1.41 | -6.90E+00 | 19 | 1.40E-06 | **** | [-14.66, -7.84] | 2.140000 | 1.31E+04 | 1.00 |
| | A1 | 37.50 ± 29.17 | 30.62 ± 20.19 | 4.67 | 3.23 | 4.21E+00 | 39 | 1.43E-04 | *** | [3.58, 10.17] | 0.270000 | 1.75E+02 | 0.39 |
| | mTVA | 45.42 ± 9.67 | 47.29 ± 6.29 | 1.55 | 1.01 | -9.50E-01 | 39 | 3.47E-01 | ns | [-5.86, 2.11] | 0.230000 | 2.60E-01 | 0.29 |
| | pTVA | 65.00 ± 7.26 | 54.58 ± 4.15 | 1.16 | 0.66 | 9.78E+00 | 39 | 4.83E-12 | **** | [8.26, 12.57] | 1.740000 | 1.83E+09 | 1.00 |
| | aTVA | 60.42 ± 5.19 | 63.96 ± 12.55 | 0.83 | 2.01 | -2.25E+00 | 39 | 3.03E-02 | * | [-6.73, -0.35] | 0.360000 | 1.61E+00 | 0.61 |
| | TVAs | 56.94 ± 11.30 | 55.28 ± 10.86 | 1.04 | 1.00 | 1.56E+00 | 119 | 1.22E-01 | ns | [-0.45, 3.78] | 0.150000 | 3.28E-01 | 0.37 |
| Identity | LA1 | 0.15 ± 0.64 | 0.29 ± 1.28 | 0.15 | 0.29 | -4.40E-01 | 19 | 6.66E-01 | ns | [-0.85, 0.56] | 0.140000 | 2.53E-01 | 0.09 |
| | RA1 | 11.47 ± 5.09 | 18.09 ± 3.26 | 1.17 | 0.75 | -4.58E+00 | 19 | 2.05E-04 | *** | [-9.64, -3.59] | 1.510000 | 1.47E+02 | 1.00 |
| | LmTVA | 11.47 ± 4.73 | 11.18 ± 4.01 | 1.09 | 0.92 | 2.10E-01 | 19 | 8.39E-01 | ns | [-2.70, 3.29] | 0.070000 | 2.37E-01 | 0.06 |
| | RmTVA | 0.59 ± 1.50 | 2.35 ± 3.03 | 0.34 | 0.69 | -2.11E+00 | 19 | 4.86E-02 | * | [-3.52, -0.01] | 0.720000 | 1.42E+00 | 0.86 |
| | LpTVA | 9.71 ± 3.37 | 12.21 ± 3.39 | 0.77 | 0.78 | -2.43E+00 | 19 | 2.53E-02 | * | [-4.65, -0.35] | 0.720000 | 2.39E+00 | 0.86 |
| | RpTVA | 22.65 ± 2.10 | 6.76 ± 4.66 | 0.48 | 1.07 | 1.31E+01 | 19 | 5.99E-11 | **** | [13.34, 18.42] | 4.280000 | 1.48E+08 | 1.00 |
| | LaTVA | 10.29 ± 4.51 | 11.47 ± 1.28 | 1.03 | 0.29 | -1.00E+00 | 19 | 3.30E-01 | ns | [-3.64, 1.29] | 0.350000 | 3.61E-01 | 0.31 |
| | RaTVA | 6.18 ± 3.94 | 7.35 ± 8.29 | 0.90 | 1.90 | -7.50E-01 | 19 | 4.64E-01 | ns | [-4.47, 2.12] | 0.180000 | 2.98E-01 | 0.12 |
| | A1 | 5.81 ± 6.72 | 9.19 ± 9.24 | 1.08 | 1.48 | -3.77E+00 | 39 | 5.40E-04 | *** | [-5.20, -1.57] | 0.410000 | 5.30E+01 | 0.72 |
| | mTVA | 6.03 ± 6.48 | 6.76 ± 5.67 | 1.04 | 0.91 | -8.80E-01 | 39 | 3.83E-01 | ns | [-2.42, 0.95] | 0.120000 | 2.45E-01 | 0.11 |
| | pTVA | 16.18 ± 7.05 | 9.49 ± 4.90 | 1.13 | 0.78 | 4.01E+00 | 39 | 2.65E-04 | *** | [3.32, 10.07] | 1.090000 | 1.00E+02 | 1.00 |
| | aTVA | 8.24 ± 4.71 | 9.41 ± 6.28 | 0.75 | 1.01 | -1.21E+00 | 39 | 2.32E-01 | ns | [-3.14, 0.79] | 0.210000 | 3.37E-01 | 0.25 |
| | TVAs | 10.15 ± 7.55 | 8.55 ± 5.78 | 0.69 | 0.53 | 2.07E+00 | 119 | 4.06E-02 | * | [0.07, 3.12] | 0.240000 | 7.94E-01 | 0.73 |

**Supplementary Table 11. Comparing the performance of the models decoding speaker identity-related information.** This table reports the significance of the speaker identity decoding VLS-LIN difference. Paired t-tests were conducted between the mean scores of linear classifiers pre-trained to detect gender (2 classes), age (2 classes), and identity (17 classes) from the VLS features, and those trained with the LIN features. These scores were obtained after classifying the VLS or LIN coordinates of the 18 Test Stimuli (3 participants x 6 stimuli per participant). s.e.m. = standard error of the mean. Here are reported the results of the statistical tests, t-value, degree of freedom (dof), p-value, and degree of significance (unc. sig.), 95% confidence interval (CI95%), effect size (Cohen-d), Bayes Factor (BF10), and statistical power (power) for each speaker information and ROI.

| Category | Model | ROI | Accuracy ROI (%) | Accuracy A1 (%) | s.e.m. ROI | s.e.m. A1 | T ROI vs A1 | dof | p-val | unc. sig. | CI95% | cohen-d | BF10 | power |
|---|---|---|---|---|---|---|---|---|---|---|---|---|---|---|
| Gender | LIN | mTVA | 50.14 ± 11.28 | 47.08 ± 4.30 | 1.81 | 0.69 | 1.58E+00 | 78 | 1.20E-01 | ns | [-0.79, 6.90] | 3.50E-01 | 6.83E-01 | 0.35 |
| | | pTVA | 72.08 ± 5.48 | 47.08 ± 4.30 | 0.88 | 0.69 | 2.24E+01 | 78 | 0.00E+00 | **** | [22.78, 27.22] | 5.01E+00 | 1.30E+32 | 1.00 |
| | | aTVA | 44.44 ± 0.00 | 47.08 ± 4.30 | 0.00 | 0.69 | -3.83E+00 | 78 | 0.00E+00 | *** | [-4.01, -1.27] | 8.60E-01 | 9.78E+01 | 0.97 |
| | | TVAs | 55.56 ± 13.94 | 47.08 ± 4.30 | 1.28 | 0.69 | 3.76E+00 | 158 | 0.00E+00 | *** | [4.02, 12.92] | 6.90E-01 | 9.90E+01 | 0.96 |
| | VLS | mTVA | 50.00 ± 5.56 | 61.11 ± 2.15 | 0.89 | 0.34 | -1.17E+01 | 78 | 0.00E+00 | **** | [-13.01, -9.21] | 2.60E+00 | 3.45E+15 | 1.00 |
| | | pTVA | 63.89 ± 2.78 | 61.11 ± 2.15 | 0.44 | 0.34 | 4.94E+00 | 78 | 0.00E+00 | **** | [1.66, 3.90] | 1.10E+00 | 3.59E+03 | 1.00 |
| | | aTVA | 47.50 ± 3.72 | 61.11 ± 2.15 | 0.60 | 0.34 | -1.98E+01 | 78 | 0.00E+00 | **** | [-14.98, -12.24] | 4.43E+00 | 4.06E+28 | 1.00 |
| | | TVAs | 53.80 ± 8.33 | 61.11 ± 2.15 | 0.76 | 0.34 | -5.46E+00 | 158 | 0.00E+00 | **** | [-9.96, -4.67] | 1.00E+00 | 6.55E+04 | 1.00 |
| Age | LIN | mTVA | 47.29 ± 6.29 | 30.62 ± 20.19 | 1.01 | 3.23 | 4.92E+00 | 78 | 0.00E+00 | **** | [9.93, 23.41] | 1.10E+00 | 3.41E+03 | 1.00 |
| | | pTVA | 54.58 ± 4.15 | 30.62 ± 20.19 | 0.66 | 3.23 | 7.26E+00 | 78 | 0.00E+00 | **** | [17.39, 30.53] | 1.62E+00 | 3.02E+07 | 1.00 |
| | | aTVA | 63.96 ± 12.55 | 30.62 ± 20.19 | 2.01 | 3.23 | 8.76E+00 | 78 | 0.00E+00 | **** | [25.75, 40.91] | 1.96E+00 | 1.68E+10 | 1.00 |
| | | TVAs | 55.28 ± 10.86 | 30.62 ± 20.19 | 1.00 | 3.23 | 9.72E+00 | 158 | 0.00E+00 | **** | [19.65, 29.66] | 1.78E+00 | 4.55E+14 | 1.00 |
| | VLS | mTVA | 45.42 ± 9.67 | 37.50 ± 29.17 | 1.55 | 4.67 | 1.61E+00 | 78 | 1.10E-01 | ns | [-1.88, 17.71] | 3.60E-01 | 7.10E-01 | 0.36 |
| | | pTVA | 65.00 ± 7.26 | 37.50 ± 29.17 | 1.16 | 4.67 | 5.71E+00 | 78 | 0.00E+00 | **** | [17.92, 37.08] | 1.28E+00 | 6.21E+04 | 1.00 |
| | | aTVA | 60.42 ± 5.19 | 37.50 ± 29.17 | 0.83 | 4.67 | 4.83E+00 | 78 | 0.00E+00 | **** | [13.47, 32.36] | 1.08E+00 | 2.48E+03 | 1.00 |
| | | TVAs | 56.94 ± 11.30 | 37.50 ± 29.17 | 1.04 | 4.67 | 6.03E+00 | 158 | 0.00E+00 | **** | [13.07, 25.82] | 1.10E+00 | 8.68E+05 | 1.00 |
| Identity | LIN | mTVA | 6.76 ± 5.67 | 9.19 ± 9.24 | 0.91 | 1.48 | -1.40E+00 | 78 | 1.70E-01 | ns | [-5.88, 1.03] | 3.10E-01 | 5.42E-01 | 0.28 |
| | | pTVA | 9.49 ± 4.90 | 9.19 ± 9.24 | 0.78 | 1.48 | 1.80E-01 | 78 | 8.60E-01 | ns | [-3.04, 3.63] | 4.00E-02 | 2.36E-01 | 0.05 |
| | | aTVA | 9.41 ± 6.28 | 9.19 ± 9.24 | 1.01 | 1.48 | 1.20E-01 | 78 | 9.00E-01 | ns | [-3.34, 3.78] | 3.00E-02 | 2.34E-01 | 0.05 |
| | | TVAs | 8.55 ± 5.78 | 9.19 ± 9.24 | 0.53 | 1.48 | -5.10E-01 | 158 | 6.10E-01 | ns | [-3.11, 1.83] | 9.00E-02 | 2.19E-01 | 0.08 |
| | VLS | mTVA | 6.03 ± 6.48 | 5.81 ± 6.72 | 1.04 | 1.08 | 1.50E-01 | 78 | 8.80E-01 | ns | [-2.76, 3.20] | 3.00E-02 | 2.35E-01 | 0.05 |
| | | pTVA | 16.18 ± 7.05 | 5.81 ± 6.72 | 1.13 | 1.08 | 6.65E+00 | 78 | 0.00E+00 | **** | [7.26, 13.47] | 1.49E+00 | 2.43E+06 | 1.00 |
| | | aTVA | 8.24 ± 4.71 | 5.81 ± 6.72 | 0.75 | 1.08 | 1.85E+00 | 78 | 7.00E-02 | ns | [-0.19, 5.04] | 4.10E-01 | 1.01E+00 | 0.45 |
| | | TVAs | 10.15 ± 7.55 | 5.81 ± 6.72 | 0.69 | 1.08 | 3.21E+00 | 158 | 0.00E+00 | ** | [1.67, 7.00] | 5.90E-01 | 1.91E+01 | 0.89 |

**Supplementary Table 12. Comparing the performance of the models decoding speaker identity-related information by ROI.** This table reports the significance of the speaker identity decoding A1-TVAs difference. Two-sample t-tests were conducted for each model to determine if there was an A1-TVAs difference between the mean scores of linear classifiers pre-trained to detect gender (2 classes), age (2 classes), and identity (17 classes). These scores were obtained by classifying the VLS coordinates or LIN features, reconstructed by different ROIs, for the 18 Test Stimuli (3 participants x 6 stimuli per participant). s.e.m. = standard error of the mean. Here are reported the results of the statistical tests, t-value, degree of freedom (dof), p-value, degree of significance (unc. sig.), 95% confidence interval (CI95%), effect size (Cohen-d), Bayes Factor (BF10), and statistical power (power) for each speaker information and model.

| Model | ROI | Accuracy (%) | s.e.m. | T | dof | p-val | unc. sig. | CI95% | cohen-d | BF10 | power |
|---|---|---|---|---|---|---|---|---|---|---|---|
| LIN | LA1 | 44.87 ± 7.99 | 2.31 | n/a | n/a | n/a | n/a | n/a | n/a | n/a | n/a |
| | RA1 | 51.28 ± 10.02 | 2.89 | 4.40E-01 | 12 | 3.33E-01 | ns | [46.13, inf] | 1.20E-01 | 6.06E-01 | 0.11 |
| | LmTVA | 51.71 ± 10.76 | 3.11 | 5.50E-01 | 12 | 2.96E-01 | ns | [46.17, inf] | 1.50E-01 | 6.35E-01 | 0.13 |
| | RmTVA | 43.59 ± 8.40 | 2.42 | n/a | n/a | n/a | n/a | n/a | n/a | n/a | n/a |
| | LpTVA | 50.00 ± 8.72 | 2.52 | n/a | n/a | n/a | n/a | n/a | n/a | n/a | n/a |
| | RpTVA | 52.99 ± 10.36 | 2.99 | 1.00E+00 | 12 | 1.69E-01 | ns | [47.66, inf] | 2.80E-01 | 8.49E-01 | 0.24 |
| | LaTVA | 51.28 ± 8.48 | 2.45 | 5.20E-01 | 12 | 3.05E-01 | ns | [46.92, inf] | 1.50E-01 | 6.27E-01 | 0.13 |
| | RaTVA | 45.30 ± 9.71 | 2.80 | n/a | n/a | n/a | n/a | n/a | n/a | n/a | n/a |
| | A1 | 48.08 ± 9.62 | 1.92 | n/a | n/a | n/a | n/a | n/a | n/a | n/a | n/a |
| | mTVA | 47.65 ± 10.47 | 2.09 | n/a | n/a | n/a | n/a | n/a | n/a | n/a | n/a |
| | pTVA | 51.50 ± 9.69 | 1.94 | 7.70E-01 | 25 | 2.24E-01 | ns | [48.18, inf] | 1.50E-01 | 5.43E-01 | 0.19 |
| | aTVA | 48.29 ± 9.59 | 1.92 | n/a | n/a | n/a | n/a | n/a | n/a | n/a | n/a |
| | TVAs | 49.15 ± 10.07 | 1.15 | n/a | n/a | n/a | n/a | n/a | n/a | n/a | n/a |
| VLS | LA1 | 50.00 ± 11.32 | 3.27 | n/a | n/a | n/a | n/a | n/a | n/a | n/a | n/a |
| | RA1 | 61.54 ± 6.34 | 1.83 | 6.31E+00 | 12 | 1.96E-05 | **** | [58.28, inf] | 1.75E+00 | 1.31E+03 | 1.00 |
| | LmTVA | 51.71 ± 5.06 | 1.46 | 1.17E+00 | 12 | 1.32E-01 | ns | [49.11, inf] | 3.20E-01 | 9.85E-01 | 0.29 |
| | RmTVA | 45.73 ± 8.76 | 2.53 | n/a | n/a | n/a | n/a | n/a | n/a | n/a | n/a |
| | LpTVA | 63.25 ± 7.40 | 2.14 | 6.20E+00 | 12 | 2.29E-05 | **** | [59.44, inf] | 1.72E+00 | 1.14E+03 | 1.00 |
| | RpTVA | 60.26 ± 6.48 | 1.87 | 5.48E+00 | 12 | 7.01E-05 | **** | [56.92, inf] | 1.52E+00 | 4.30E+02 | 1.00 |
| | LaTVA | 60.26 ± 6.10 | 1.76 | 5.82E+00 | 12 | 4.10E-05 | **** | [57.12, inf] | 1.61E+00 | 6.86E+02 | 1.00 |
| | RaTVA | 50.00 ± 8.98 | 2.59 | n/a | n/a | n/a | n/a | n/a | n/a | n/a | n/a |
| | A1 | 55.77 ± 10.84 | 2.17 | 2.66E+00 | 25 | 6.70E-03 | ** | [52.07, inf] | 5.20E-01 | 7.39E+00 | 0.83 |
| | mTVA | 48.72 ± 7.75 | 1.55 | n/a | n/a | n/a | n/a | n/a | n/a | n/a | n/a |
| | pTVA | 61.75 ± 7.12 | 1.42 | 8.26E+00 | 25 | 6.56E-09 | **** | [59.32, inf] | 1.62E+00 | 2.00E+06 | 1.00 |
| | aTVA | 55.13 ± 9.24 | 1.85 | 2.78E+00 | 25 | 5.13E-03 | ** | [51.97, inf] | 5.40E-01 | 9.24E+00 | 0.85 |
| | TVAs | 55.20 ± 9.68 | 1.10 | 4.71E+00 | 77 | 5.29E-06 | **** | [53.36, inf] | 5.30E-01 | 3.23E+03 | 1.00 |

**Supplementary Table 13. Assessing the significance of the speaker gender categorization task.** This table reports the significance of the speaker's gender categorization performance. 342 voice stimuli were used in the experiments: the original stimuli (N = 18), directly reconstructed stimuli using the LIN and the VLS models (N = 36), and brain-reconstructed stimuli (18 stimuli x 2 models x 4 regions of interest x 2 hemispheres, N = 288). The participants were tasked with identifying the gender of the presented voice in each trial by clicking either the ‘Female’ or ‘Male’ button. To evaluate the accuracy of the binary responses, we computed the classification accuracy for each participant and region of interest

(ROI). We then utilized one-sample t-tests to compare the mean accuracy distribution across all participants to the chance level of 50%. s.e.m. = standard error of the mean. Here are reported the results of the statistical tests, t-value, degree of freedom (dof), p-value, degree of significance (unc. sig.), 95% confidence interval (CI95%), effect size (Cohen-d), Bayes Factor (BF10), and statistical power (power) for each model and ROI.

| Model | ROI | Accuracy (%) | s.e.m. | T | dof | p-val | unc. sig. | CI95% | cohen-d | BF10 | power |
|---|---|---|---|---|---|---|---|---|---|---|---|
| LIN | LA1 | 46.15 ± 14.48 | 4.18 | n/a | n/a | n/a | n/a | n/a | n/a | n/a | n/a |
| | RA1 | 44.23 ± 11.50 | 3.32 | n/a | n/a | n/a | n/a | n/a | n/a | n/a | n/a |
| | LmTVA | 50.00 ± 9.81 | 2.83 | n/a | n/a | n/a | n/a | n/a | n/a | n/a | n/a |
| | RmTVA | 57.69 ± 12.85 | 3.71 | 2.07E+00 | 12 | 3.02E-02 | * | [51.08, inf] | 5.80E-01 | 2.80E+00 | 0.62 |
| | LpTVA | 50.00 ± 10.34 | 2.98 | 0.00E+00 | 12 | 5.00E-01 | ns | [44.68, inf] | 0.00E+00 | 5.56E-01 | 0.05 |
| | RpTVA | 50.64 ± 10.57 | 3.05 | 2.10E-01 | 12 | 4.19E-01 | ns | [45.20, inf] | 6.00E-02 | 5.67E-01 | 0.07 |
| | LaTVA | 48.72 ± 13.01 | 3.76 | n/a | n/a | n/a | n/a | n/a | n/a | n/a | n/a |
| | RaTVA | 62.82 ± 13.32 | 3.85 | 3.33E+00 | 12 | 2.98E-03 | ** | [55.97, inf] | 9.20E-01 | 1.77E+01 | 0.93 |
| | A1 | 45.19 ± 13.11 | 2.62 | n/a | n/a | n/a | n/a | n/a | n/a | n/a | n/a |
| | mTVA | 53.85 ± 12.06 | 2.41 | 1.59E+00 | 25 | 6.17E-02 | ns | [49.73, inf] | 3.10E-01 | 1.26E+00 | 0.46 |
| | pTVA | 50.32 ± 10.46 | 2.09 | 1.50E-01 | 25 | 4.40E-01 | ns | [46.75, inf] | 3.00E-02 | 4.19E-01 | 0.07 |
| | aTVA | 55.77 ± 14.94 | 2.99 | 1.93E+00 | 25 | 3.24E-02 | * | [50.67, inf] | 3.80E-01 | 2.06E+00 | 0.59 |
| | TVAs | 53.31 ± 12.82 | 1.46 | 2.27E+00 | 77 | 1.31E-02 | * | [50.88, inf] | 2.60E-01 | 2.77E+00 | 0.73 |
| VLS | LA1 | 54.49 ± 10.65 | 3.07 | 1.46E+00 | 12 | 8.50E-02 | ns | [49.01, inf] | 4.00E-01 | 1.32E+00 | 0.39 |
| | RA1 | 50.00 ± 8.01 | 2.31 | 0.00E+00 | 12 | 5.00E-01 | ns | [45.88, inf] | 0.00E+00 | 5.56E-01 | 0.05 |
| | LmTVA | 51.28 ± 10.26 | 2.96 | 4.30E-01 | 12 | 3.36E-01 | ns | [46.01, inf] | 1.20E-01 | 6.04E-01 | 0.11 |
| | RmTVA | 54.49 ± 10.65 | 3.07 | 1.46E+00 | 12 | 8.50E-02 | ns | [49.01, inf] | 4.00E-01 | 1.32E+00 | 0.39 |
| | LpTVA | 45.51 ± 11.14 | 3.22 | n/a | n/a | n/a | n/a | n/a | n/a | n/a | n/a |
| | RpTVA | 56.41 ± 8.74 | 2.52 | 2.54E+00 | 12 | 1.30E-02 | * | [51.91, inf] | 7.00E-01 | 5.38E+00 | 0.77 |
| | LaTVA | 64.74 ± 7.42 | 2.14 | 6.88E+00 | 12 | 8.46E-06 | **** | [60.93, inf] | 1.91E+00 | 2.74E+03 | 1.00 |
| | RaTVA | 61.54 ± 14.81 | 4.28 | 2.70E+00 | 12 | 9.68E-03 | ** | [53.92, inf] | 7.50E-01 | 6.79E+00 | 0.82 |
| | A1 | 52.24 ± 9.68 | 1.94 | 1.16E+00 | 25 | 1.29E-01 | ns | [48.94, inf] | 2.30E-01 | 7.57E-01 | 0.30 |
| | mTVA | 52.88 ± 10.58 | 2.12 | 1.36E+00 | 25 | 9.24E-02 | ns | [49.27, inf] | 2.70E-01 | 9.47E-01 | 0.38 |
| | pTVA | 50.96 ± 11.40 | 2.28 | 4.20E-01 | 25 | 3.38E-01 | ns | [47.07, inf] | 8.00E-02 | 4.50E-01 | 0.11 |
| | aTVA | 63.14 ± 11.82 | 2.36 | 5.56E+00 | 25 | 4.45E-06 | **** | [59.10, inf] | 1.09E+00 | 4.79E+03 | 1.00 |
| | TVAs | 55.66 ± 12.48 | 1.42 | 3.98E+00 | 77 | 7.72E-05 | **** | [53.29, inf] | 4.50E-01 | 2.69E+02 | 0.99 |

**Supplementary Table 14. Assessing the significance of the speaker age categorization task.** This table reports the significance of the speaker age categorization performance. 342 voice stimuli were used in the experiments: the original stimuli (N = 18), directly reconstructed stimuli using the LIN and the VLS models (N = 36), and brain-reconstructed stimuli (18 stimuli x 2 models x 4 regions of interest x 2 hemispheres, N = 288). The participants were tasked with identifying the approximate age of the presented voice in each trial by clicking either the 'Younger' or 'Older' button. To evaluate the accuracy of the binary responses, we computed the classification accuracy for each participant and region of interest (ROI). We then utilized one-sample t-tests to compare the mean accuracy distribution across all participants to the chance level of 50%. s.e.m. = standard error of the mean. Here are reported the results of the statistical tests, t-value, degree of freedom (dof), p-value, degree of significance (unc. sig.), 95% confidence interval (CI95%), effect size (Cohen-d), Bayes Factor (BF10), and statistical power (power) for each model and ROI.

| Model | ROI | Accuracy (%) | s.e.m. | T | dof | p-val | unc. sig. | CI95% | cohen-d | BF10 | power |
|---|---|---|---|---|---|---|---|---|---|---|---|
| LIN | LA1 | 54.70 ± 9.89 | 3.50 | 1.34E+00 | 8 | 1.08E-01 | ns | [48.20, inf] | 4.50E-01 | 1.29E+00 | 0.34 |
| | RA1 | 57.41 ± 8.69 | 3.07 | 2.41E+00 | 8 | 2.12E-02 | * | [51.70, inf] | 8.00E-01 | 4.14E+00 | 0.71 |
| | LmTVA | 57.04 ± 7.77 | 2.75 | 2.56E+00 | 8 | 1.68E-02 | * | [51.93, inf] | 8.50E-01 | 4.94E+00 | 0.76 |
| | RmTVA | 42.36 ± 8.22 | 2.91 | n/a | n/a | n/a | n/a | n/a | n/a | n/a | n/a |
| | LpTVA | 57.78 ± 7.70 | 2.72 | 2.86E+00 | 8 | 1.06E-02 | * | [52.72, inf] | 9.50E-01 | 7.04E+00 | 0.83 |
| | RpTVA | 50.98 ± 9.61 | 3.40 | 2.90E-01 | 8 | 3.90E-01 | ns | [44.67, inf] | 1.00E-01 | 6.67E-01 | 0.08 |
| | LaTVA | 40.48 ± 9.52 | 3.37 | n/a | n/a | n/a | n/a | n/a | n/a | n/a | n/a |
| | RaTVA | 40.52 ± 7.04 | 2.49 | n/a | n/a | n/a | n/a | n/a | n/a | n/a | n/a |
| | A1 | 56.05 ± 9.41 | 2.28 | 2.65E+00 | 17 | 8.36E-03 | ** | [52.09, inf] | 6.30E-01 | 6.92E+00 | 0.82 |
| | mTVA | 49.70 ± 10.85 | 2.63 | n/a | n/a | n/a | n/a | n/a | n/a | n/a | n/a |
| | pTVA | 54.38 ± 9.34 | 2.27 | 1.93E+00 | 17 | 3.51E-02 | * | [50.44, inf] | 4.60E-01 | 2.22E+00 | 0.58 |
| | aTVA | 40.50 ± 8.37 | 2.03 | n/a | n/a | n/a | n/a | n/a | n/a | n/a | n/a |
| | TVAs | 48.19 ± 11.18 | 1.54 | n/a | n/a | n/a | n/a | n/a | n/a | n/a | n/a |
| VLS | LA1 | 72.22 ± 9.16 | 3.24 | 6.86E+00 | 8 | 6.48E-05 | **** | [66.20, inf] | 2.29E+00 | 4.52E+02 | 1.00 |
| | RA1 | 48.33 ± 5.77 | 2.04 | n/a | n/a | n/a | n/a | n/a | n/a | n/a | n/a |
| | LmTVA | 51.46 ± 7.76 | 2.74 | 5.30E-01 | 8 | 3.04E-01 | ns | [46.36, inf] | 1.80E-01 | 7.25E-01 | 0.12 |
| | RmTVA | 41.11 ± 6.57 | 2.32 | n/a | n/a | n/a | n/a | n/a | n/a | n/a | n/a |
| | LpTVA | 60.61 ± 5.67 | 2.00 | 5.29E+00 | 8 | 3.68E-04 | *** | [56.88, inf] | 1.76E+00 | 1.06E+02 | 1.00 |
| | RpTVA | 66.05 ± 6.65 | 2.35 | 6.83E+00 | 8 | 6.70E-05 | **** | [61.68, inf] | 2.28E+00 | 4.40E+02 | 1.00 |
| | LaTVA | 52.02 ± 8.33 | 2.94 | 6.90E-01 | 8 | 2.56E-01 | ns | [46.54, inf] | 2.30E-01 | 7.83E-01 | 0.15 |
| | RaTVA | 50.00 ± 7.53 | 2.66 | n/a | n/a | n/a | n/a | n/a | n/a | n/a | n/a |
| | A1 | 60.28 ± 14.19 | 3.44 | 2.99E+00 | 17 | 4.14E-03 | ** | [54.29, inf] | 7.00E-01 | 1.24E+01 | 0.89 |
| | mTVA | 46.29 ± 8.86 | 2.15 | n/a | n/a | n/a | n/a | n/a | n/a | n/a | n/a |
| | pTVA | 63.33 ± 6.75 | 1.64 | 8.14E+00 | 17 | 1.44E-07 | **** | [60.48, inf] | 1.92E+00 | 1.11E+05 | 1.00 |
| | aTVA | 51.01 ± 8.00 | 1.94 | 5.20E-01 | 17 | 3.05E-01 | ns | [47.63, inf] | 1.20E-01 | 5.49E-01 | 0.13 |
| | TVAs | 53.54 ± 10.69 | 1.47 | 2.41E+00 | 53 | 9.69E-03 | ** | [51.08, inf] | 3.30E-01 | 4.15E+00 | 0.77 |

**Supplementary Table 15. Assessing the significance of the speaker identity discrimination task.** This table reports the significance of the speaker identity discrimination performance. The participants listened to 684 voice stimuli with short breaks in between. Each trial contained 2 short sound samples, and the participants had to indicate whether the samples were from the same speaker or different speakers. We then utilized one-sample t-tests to compare the mean accuracy distribution across all participants to the chance level of 50%. s.e.m. = standard error of the mean. Here are reported the results of the statistical tests, t-value, degree of freedom (dof), p-value, degree of significance (unc. sig.), 95% confidence interval (CI95%), effect size (Cohen-d), Bayes Factor (BF10), and statistical power (power) for each model and ROI.

| Category | ROI | Accuracy VLS (%) | Accuracy LIN (%) | s.e.m. VLS | s.e.m. LIN | T VLS vs LIN | dof | p-val | unc. sig. | CI95% | cohen-d | BF10 | power |
|---|---|---|---|---|---|---|---|---|---|---|---|---|---|
| Gender | LA1 | 48.33 ± 10.56 | 46.11 ± 6.60 | 3.52 | 2.20 | 4.70E-01 | 9 | 6.48E-01 | ns | [-8.41, 12.85] | 0.240000 | 3.40E-01 | 0.10 |
| | RA1 | 60.00 ± 5.98 | 53.33 ± 8.31 | 1.99 | 2.77 | 1.86E+00 | 9 | 9.63E-02 | ns | [-1.46, 14.79] | 0.870000 | 1.08E+00 | 0.69 |
| | LmTVA | 51.67 ± 5.58 | 50.56 ± 10.08 | 1.86 | 3.36 | 4.10E-01 | 9 | 6.93E-01 | ns | [-5.05, 7.27] | 0.130000 | 3.32E-01 | 0.07 |
| | RmTVA | 46.67 ± 9.03 | 43.33 ± 8.53 | 3.01 | 2.84 | 1.20E+00 | 9 | 2.60E-01 | ns | [-2.94, 9.60] | 0.360000 | 5.51E-01 | 0.18 |
| | LpTVA | 63.89 ± 7.95 | 49.44 ± 9.44 | 2.65 | 3.15 | 3.03E+00 | 9 | 1.43E-02 | * | [3.65, 25.24] | 1.570000 | 4.66E+00 | 0.99 |
| | RpTVA | 62.22 ± 4.84 | 53.33 ± 10.60 | 1.61 | 3.53 | 1.95E+00 | 9 | 8.26E-02 | ns | [-1.41, 19.18] | 1.020000 | 1.21E+00 | 0.82 |
| | LaTVA | 60.00 ± 5.44 | 50.56 ± 8.77 | 1.81 | 2.92 | 2.68E+00 | 9 | 2.50E-02 | * | [1.49, 17.40] | 1.230000 | 3.00E+00 | 0.93 |
| | RaTVA | 50.00 ± 9.94 | 45.56 ± 9.88 | 3.31 | 3.29 | 1.15E+00 | 9 | 2.80E-01 | ns | [-4.30, 13.19] | 0.430000 | 5.26E-01 | 0.23 |
| | A1 | 54.17 ± 10.37 | 49.72 ± 8.33 | 2.38 | 1.91 | 1.52E+00 | 19 | 1.45E-01 | ns | [-1.67, 10.56] | 0.460000 | 6.25E-01 | 0.50 |
| | mTVA | 49.17 ± 7.91 | 46.94 ± 10.01 | 1.81 | 2.30 | 1.16E+00 | 19 | 2.58E-01 | ns | [-1.77, 6.21] | 0.240000 | 4.21E-01 | 0.18 |
| | pTVA | 63.06 ± 6.64 | 51.39 ± 10.23 | 1.52 | 2.35 | 3.57E+00 | 19 | 2.06E-03 | ** | [4.82, 18.51] | 1.320000 | 1.95E+01 | 1.00 |
| | aTVA | 55.00 ± 9.44 | 48.06 ± 9.67 | 2.17 | 2.22 | 2.66E+00 | 19 | 1.54E-02 | * | [1.49, 12.40] | 0.710000 | 3.59E+00 | 0.85 |
| | TVAs | 55.74 ± 9.88 | 48.80 ± 10.15 | 1.29 | 1.32 | 4.37E+00 | 59 | 5.05E-05 | **** | [3.77, 10.12] | 0.690000 | 4.08E+02 | 1.00 |
| Age | LA1 | 54.49 ± 10.65 | 46.15 ± 14.48 | 3.07 | 4.18 | 1.54E+00 | 12 | 1.50E-01 | ns | [-3.48, 20.14] | 0.630000 | 7.18E-01 | 0.55 |
| | RA1 | 50.00 ± 8.01 | 44.23 ± 11.50 | 2.31 | 3.32 | 1.24E+00 | 12 | 2.39E-01 | ns | [-4.38, 15.92] | 0.560000 | 5.25E-01 | 0.46 |
| | LmTVA | 51.28 ± 10.26 | 50.00 ± 9.81 | 2.96 | 2.83 | 2.70E-01 | 12 | 7.94E-01 | ns | [-9.17, 11.73] | 0.120000 | 2.87E-01 | 0.07 |
| | RmTVA | 54.49 ± 10.65 | 57.69 ± 12.85 | 3.07 | 3.71 | -7.20E-01 | 12 | 4.88E-01 | ns | [-12.97, 6.55] | 0.260000 | 3.47E-01 | 0.14 |
| | LpTVA | 45.51 ± 11.14 | 50.00 ± 10.34 | 3.22 | 2.98 | -1.17E+00 | 12 | 2.66E-01 | ns | [-12.87, 3.89] | 0.400000 | 4.90E-01 | 0.27 |
| | RpTVA | 56.41 ± 8.74 | 50.64 ± 10.57 | 2.52 | 3.05 | 1.74E+00 | 12 | 1.08E-01 | ns | [-1.47, 13.00] | 0.570000 | 9.09E-01 | 0.47 |
| | LaTVA | 64.74 ± 7.42 | 48.72 ± 13.01 | 2.14 | 3.76 | 5.25E+00 | 12 | 2.04E-04 | *** | [9.38, 22.68] | 1.450000 | 1.55E+02 | 1.00 |
| | RaTVA | 61.54 ± 14.81 | 62.82 ± 13.32 | 4.28 | 3.85 | -2.50E-01 | 12 | 8.08E-01 | ns | [-12.51, 9.95] | 0.090000 | 2.86E-01 | 0.06 |
| | A1 | 52.24 ± 9.68 | 45.19 ± 13.11 | 1.94 | 2.62 | 2.01E+00 | 25 | 5.55E-02 | ns | [-0.18, 14.28] | 0.600000 | 1.16E+00 | 0.84 |
| | mTVA | 52.88 ± 10.58 | 53.85 ± 12.06 | 2.12 | 2.41 | -3.00E-01 | 25 | 7.70E-01 | ns | [-7.65, 5.72] | 0.080000 | 2.16E-01 | 0.07 |
| | pTVA | 50.96 ± 11.40 | 50.32 ± 10.46 | 2.28 | 2.09 | 2.40E-01 | 25 | 8.14E-01 | ns | [-4.90, 6.19] | 0.060000 | 2.13E-01 | 0.06 |
| | aTVA | 63.14 ± 11.82 | 55.77 ± 14.94 | 2.36 | 2.99 | 2.16E+00 | 25 | 4.02E-02 | * | [0.35, 14.39] | 0.540000 | 1.50E+00 | 0.75 |
| | TVAs | 55.66 ± 12.48 | 53.31 ± 12.82 | 1.42 | 1.46 | 1.28E+00 | 77 | 2.03E-01 | ns | [-1.29, 6.00] | 0.180000 | 2.74E-01 | 0.36 |
| Identity | LA1 | 72.22 ± 9.16 | 54.70 ± 9.89 | 3.24 | 3.50 | 3.64E+00 | 8 | 6.61E-03 | ** | [6.41, 28.63] | 1.730000 | 8.84E+00 | 0.99 |
| | RA1 | 48.33 ± 5.77 | 57.41 ± 8.69 | 2.04 | 3.07 | -1.97E+00 | 8 | 8.49E-02 | ns | [-19.72, 1.57] | 1.160000 | 1.23E+00 | 0.86 |
| | LmTVA | 51.46 ± 7.76 | 57.04 ± 7.77 | 2.74 | 2.75 | -1.44E+00 | 8 | 1.87E-01 | ns | [-14.49, 3.34] | 0.680000 | 7.08E-01 | 0.43 |
| | RmTVA | 41.11 ± 6.57 | 42.36 ± 8.22 | 2.32 | 2.91 | -2.50E-01 | 8 | 8.08E-01 | ns | [-12.72, 10.22] | 0.160000 | 3.30E-01 | 0.07 |
| | LpTVA | 60.61 ± 5.67 | 57.78 ± 7.70 | 2.00 | 2.72 | 1.05E+00 | 8 | 3.23E-01 | ns | [-3.37, 9.02] | 0.390000 | 5.02E-01 | 0.18 |
| | RpTVA | 66.05 ± 6.65 | 50.98 ± 9.61 | 2.35 | 3.40 | 4.37E+00 | 8 | 2.39E-03 | ** | [7.11, 23.03] | 1.720000 | 2.01E+01 | 0.99 |
| | LaTVA | 52.02 ± 8.33 | 40.48 ± 9.52 | 2.94 | 3.37 | 2.00E+00 | 8 | 8.02E-02 | ns | [-1.75, 24.84] | 1.220000 | 1.29E+00 | 0.89 |
| | RaTVA | 50.00 ± 7.53 | 40.52 ± 7.04 | 2.66 | 2.49 | 2.60E+00 | 8 | 3.16E-02 | * | [1.07, 17.88] | 1.230000 | 2.59E+00 | 0.89 |
| | A1 | 60.28 ± 14.19 | 56.05 ± 9.41 | 3.44 | 2.28 | 9.20E-01 | 17 | 3.68E-01 | ns | [-5.42, 13.86] | 0.340000 | 3.54E-01 | 0.28 |
| | mTVA | 46.29 ± 8.86 | 49.70 ± 10.85 | 2.15 | 2.63 | -1.10E+00 | 17 | 2.86E-01 | ns | [-9.96, 3.13] | 0.330000 | 4.11E-01 | 0.27 |
| | pTVA | 63.33 ± 6.75 | 54.38 ± 9.34 | 1.64 | 2.27 | 3.46E+00 | 17 | 3.02E-03 | ** | [3.49, 14.41] | 1.070000 | 1.45E+01 | 0.99 |
| | aTVA | 51.01 ± 8.00 | 40.50 ± 8.37 | 1.94 | 2.03 | 3.17E+00 | 17 | 5.62E-03 | ** | [3.51, 17.51] | 1.250000 | 8.55E+00 | 1.00 |
| | TVAs | 53.54 ± 10.69 | 48.19 ± 11.18 | 1.47 | 1.54 | 2.80E+00 | 53 | 7.15E-03 | ** | [1.51, 9.18] | 0.480000 | 4.88E+00 | 0.94 |

**Supplementary Table 16. Comparing human listeners' performance in discriminating speaker identity-related information decoded with VLS versus LIN.** This table reports the significance of the VLS-LIN difference in the speaker identity categorization and discrimination performance. Paired t-tests were conducted between the scores of human listeners at discriminating the speaker gender (2 classes), age (2 classes), and identity (17 classes) of the 18 Test Stimuli reconstructed from the VLS features with those from LIN features. s.e.m. = standard error of the mean. Here are reported the results of the statistical tests, t-value, degree of freedom (dof), p-value, degree of significance (unc. sig.), 95% confidence interval (CI95%), effect size (Cohen-d), Bayes Factor (BF10), and statistical power (power) for each speaker identity information and ROI.

| Category | Model | ROI | Accuracy ROI (%) | Accuracy A1 (%) | s.e.m. ROI | s.e.m. A1 | T ROI vs A1 | dof | p-val | unc. sig. | CI95% | cohen-d | BF10 | power |
|---|---|---|---|---|---|---|---|---|---|---|---|---|---|---|
| Gender | LIN | mTVA | 46.94 ± 10.01 | 49.72 ± 8.33 | 2.30 | 1.91 | -9.30E-01 | 38 | 3.60E-01 | ns | [-8.83, 3.27] | 2.90E-01 | 4.35E-01 | 0.15 |
| | | pTVA | 51.39 ± 10.23 | 49.72 ± 8.33 | 2.35 | 1.91 | 5.50E-01 | 38 | 5.80E-01 | ns | [-4.46, 7.79] | 1.70E-01 | 3.48E-01 | 0.08 |
| | | aTVA | 48.06 ± 9.67 | 49.72 ± 8.33 | 2.22 | 1.91 | -5.70E-01 | 38 | 5.70E-01 | ns | [-7.59, 4.26] | 1.80E-01 | 3.51E-01 | 0.09 |
| | | TVAs | 48.80 ± 10.15 | 49.72 ± 8.33 | 1.32 | 1.91 | -3.60E-01 | 78 | 7.20E-01 | ns | [-5.99, 4.14] | 9.00E-02 | 2.77E-01 | 0.06 |
| | VLS | mTVA | 49.17 ± 7.91 | 54.17 ± 10.37 | 1.81 | 2.38 | -1.67E+00 | 38 | 1.00E-01 | ns | [-11.06, 1.06] | 5.30E-01 | 9.19E-01 | 0.37 |
| | | pTVA | 63.06 ± 6.64 | 54.17 ± 10.37 | 1.52 | 2.38 | 3.15E+00 | 38 | 0.00E+00 | ** | [3.17, 14.61] | 9.90E-01 | 1.21E+01 | 0.87 |
| | | aTVA | 55.00 ± 9.44 | 54.17 ± 10.37 | 2.17 | 2.38 | 2.60E-01 | 38 | 8.00E-01 | ns | [-5.68, 7.35] | 8.00E-02 | 3.17E-01 | 0.06 |
| | | TVAs | 55.74 ± 9.88 | 54.17 ± 10.37 | 1.29 | 2.38 | 6.00E-01 | 78 | 5.50E-01 | ns | [-3.64, 6.78] | 1.60E-01 | 3.05E-01 | 0.09 |
| Age | LIN | mTVA | 53.85 ± 12.06 | 45.19 ± 13.11 | 2.41 | 2.62 | 2.43E+00 | 50 | 2.00E-02 | * | [1.50, 15.81] | 6.70E-01 | 2.95E+00 | 0.66 |
| | | pTVA | 50.32 ± 10.46 | 45.19 ± 13.11 | 2.09 | 2.62 | 1.53E+00 | 50 | 1.30E-01 | ns | [-1.61, 11.87] | 4.20E-01 | 7.23E-01 | 0.32 |
| | | aTVA | 55.77 ± 14.94 | 45.19 ± 13.11 | 2.99 | 2.62 | 2.66E+00 | 50 | 1.00E-02 | * | [2.59, 18.56] | 7.40E-01 | 4.65E+00 | 0.74 |
| | | TVAs | 53.31 ± 12.82 | 45.19 ± 13.11 | 1.46 | 2.62 | 2.75E+00 | 102 | 1.00E-02 | ** | [2.27, 13.97] | 6.20E-01 | 5.95E+00 | 0.78 |
| | VLS | mTVA | 52.88 ± 10.58 | 52.24 ± 9.68 | 2.12 | 1.94 | 2.20E-01 | 50 | 8.20E-01 | ns | [-5.12, 6.40] | 6.00E-02 | 2.84E-01 | 0.06 |
| | | pTVA | 50.96 ± 11.40 | 52.24 ± 9.68 | 2.28 | 1.94 | -4.30E-01 | 50 | 6.70E-01 | ns | [-7.29, 4.73] | 1.20E-01 | 3.00E-01 | 0.07 |
| | | aTVA | 63.14 ± 11.82 | 52.24 ± 9.68 | 2.36 | 1.94 | 3.56E+00 | 50 | 0.00E+00 | *** | [4.76, 17.04] | 9.90E-01 | 3.70E+01 | 0.94 |
| | | TVAs | 55.66 ± 12.48 | 52.24 ± 9.68 | 1.42 | 1.94 | 1.26E+00 | 102 | 2.10E-01 | ns | [-1.95, 8.79] | 2.90E-01 | 4.67E-01 | 0.24 |
| Identity | LIN | mTVA | 49.70 ± 10.85 | 56.05 ± 9.41 | 2.63 | 2.28 | -1.82E+00 | 34 | 8.00E-02 | ns | [-13.43, 0.72] | 6.10E-01 | 1.15E+00 | 0.43 |
| | | pTVA | 54.38 ± 9.34 | 56.05 ± 9.41 | 2.27 | 2.28 | -5.20E-01 | 34 | 6.10E-01 | ns | [-8.21, 4.86] | 1.70E-01 | 3.58E-01 | 0.08 |
| | | aTVA | 40.50 ± 8.37 | 56.05 ± 9.41 | 2.03 | 2.28 | -5.09E+00 | 34 | 0.00E+00 | **** | [-21.76, -9.35] | 1.70E+00 | 1.25E+03 | 1.00 |
| | | TVAs | 48.19 ± 11.18 | 56.05 ± 9.41 | 1.54 | 2.28 | -2.65E+00 | 70 | 1.00E-02 | * | [-13.79, -1.94] | 7.20E-01 | 4.69E+00 | 0.74 |
| | VLS | mTVA | 46.29 ± 8.86 | 60.28 ± 14.19 | 2.15 | 3.44 | -3.45E+00 | 34 | 0.00E+00 | ** | [-22.24, -5.75] | 1.15E+00 | 2.21E+01 | 0.92 |
| | | pTVA | 63.33 ± 6.75 | 60.28 ± 14.19 | 1.64 | 3.44 | 8.00E-01 | 34 | 4.30E-01 | ns | [-4.69, 10.79] | 2.70E-01 | 4.13E-01 | 0.12 |
| | | aTVA | 51.01 ± 8.00 | 60.28 ± 14.19 | 1.94 | 3.44 | -2.35E+00 | 34 | 2.00E-02 | * | [-17.30, -1.24] | 7.80E-01 | 2.53E+00 | 0.63 |
| | | TVAs | 53.54 ± 10.69 | 60.28 ± 14.19 | 1.47 | 3.44 | -2.09E+00 | 70 | 4.00E-02 | * | [-13.16, -0.31] | 5.70E-01 | 1.64E+00 | 0.54 |

**Supplementary Table 17. Comparing the performance of the human listeners at discriminating speaker identity-related information by ROI.** This table reports the significance of the A1-TVAs difference in the speaker identity categorization and discrimination performance. Two-sample t-tests were conducted between the scores of human listeners at discriminating the speaker gender (2 classes), age (2 classes), and identity (17 classes) of the 18 Test Stimuli that were reconstructed from the VLS features with those from LIN features. s.e.m. = standard error of the mean. Here are reported the results of the statistical tests, t-value, degree of freedom (dof), p-value, degree of significance (unc. sig.), 95% confidence interval (CI95%), effect size (Cohen-d), Bayes Factor (BF10), and statistical power (power) for each speaker identity information and ROI.

**Supplementary Audio 1. Voice latent space interpolation.**

The audio files are two original voice samples (A, B); the synthesized voice samples from the spectrograms of the autoencoder reconstructions of the original two voice samples (A', B'); the synthesized voice samples from the spectrograms of the linearly interpolated *voice latent space* (VLS; A_to_B; Fig. 1c).

A.wav: Original voice sample of a female Chinese speaker
A'.wav: Voice sample A.wav reconstructed by the autoencoder
B.wav: Original voice sample of a male French speaker
B'.wav: Voice sample B.wav reconstructed by the autoencoder
A_to_B_I*x*.wav: Reconstructed voice samples from the linear interpolation between A and B VLS, where *x* is the interpolation step (0.2, 0.4, 0.6, 0.8).

Link:
https://drive.google.com/drive/folders/1WQonOiO_FpQvj9mT3okVSasno_rck_u3?usp=sharing

**Supplementary Audio 2. Brain-based voice reconstructions.**

The audio files are reconstructed voice samples from the fMRI responses in the speakers' temporal voice areas (TVAs). These sounds were used in the quantitative and subjective voice identity tests (Fig. 4). The samples below are from a German and a Spanish speaker. The sounds are reconstructed for each speaker using 2 models: LIN and VLS.

example1_orig.wav: Original voice sample of a male German speaker
example1_VLS_RaTVA.wav: Reconstructed voice sample from fMRI activity in the right anterior temporal voice area (RaTVA) using the VLS model
example1_LIN_RaTVA.wav: Reconstructed voice sample from fMRI activity in the right anterior temporal voice area (RaTVA) using the LIN model

example1_SPEC_RaTVA.wav: Reconstructed voice sample from fMRI activity in the right anterior temporal voice area (RaTVA) using the audio spectrogram

example2_orig.wav: Original voice sample of a male Spanish speaker
example2_VLS_LmTVA.wav: Reconstructed voice sample from fMRI activity in the left middle voice area (LmTVA) using the VLS model
example2_LIN_LmTVA.wav: Reconstructed voice sample from fMRI activity in the left middle voice area (LmTVA) using the LIN model

example2_SPEC_LmTVA.wav: Reconstructed voice sample from fMRI activity in the left middle voice area (LmTVA) using the audio spectrogram

Link:
https://drive.google.com/drive/folders/1AwAV2zigRb9DxDt_xhyea8sp13Zvxcuk?usp=share_link
